# *Trichomonas vaginalis* extracellular vesicles suppress IFNε-mediated protection against host cell cytolysis

**DOI:** 10.1101/2024.10.02.616276

**Authors:** Joshua A. Kochanowsky, Emma L. Betts, Gabriel Encinas, Johnson Amoah, Sandip Kumar Mukherjee, Patricia J. Johnson

**Affiliations:** Department of Microbiology, Immunology and Molecular Genetics, University of California, Los Angeles, CA 90095; Department of Immunobiology, University of Arizona College of Medicine, Tucson, AZ 85724; Soka University of America, Aliso Viejo, CA 92656

**Keywords:** *Trichomonas vaginalis*, *Mycoplasma hominis*, Symbiosis, Interferon-epsilon, Innate immunity, Sexually Transmitted Infections, Parasites

## Abstract

*Trichomonas vaginalis* is a commonly acquired sexually transmitted infection (STI) often found in symbiosis with the intracellular bacterium *Mycoplasma hominis*, an opportunistic pathogen of the female reproductive tract associated with bacterial vaginosis. How this symbiosis affects infection outcomes, and the host cell innate immune response is still poorly understood. Here we show that *T. vaginalis* extracellular vesicles down-regulate a non-canonical type I interferon, interferon-epsilon, and suppress type I interferon responses. Transcriptomic analysis reveals that infection with *T. vaginalis* in symbiosis with *M. hominis* or *M. hominis* alone upregulates genes involved in the type I IFN response, but infection with *T. vaginalis* alone does not. Finally, we show that interferon-epsilon stimulation is protective against *T. vaginalis* cytoadherence and cytolysis of host cells and increases the ability of neutrophils to kill the parasite. These studies provide insight into the innate immune response induced by a highly prevalent STI and its bacterial symbiont.

## Main

*Trichomonas vaginalis* is an extracellular, protozoan parasite of the human urogenital tract and causes the most common non-viral sexually transmitted infection (STI)^1–3^. Infection can lead to adverse pregnancy outcomes, such as preterm delivery, low birth weight and infant death, and has been associated with infertility and an increased risk of cervical cancer^4–10^. Despite the high prevalence of infection and long-term consequences for reproductive health, the molecular pathogenesis of the disease is poorly understood. Additionally, accumulating epidemiological data indicate an unusually high prevalence of *T. vaginalis* infection in women at the age of peri-/post-menopause^11–16^. Symbiosis of the parasite with *Mycoplasma hominis,* an obligate intracellular bacterium associated with dysbiosis, bacterial vaginosis, and capable of replicating within parasite and human cells, further complicates our understanding of how *T. vaginalis* causes disease^17,18^. Although estimates of *T. vaginalis* clinical isolates that harbor *M. hominis* can exceed 85% depending on geographical location, very little is known about how this symbiosis affects *T. vaginalis* pathogenesis^19–28^.

*T. vaginalis* and other eukaryotic pathogens secrete extracellular vesicles (EVs) that contain parasite proteins, nucleic acids, and other small molecules as a form of intracellular communication between pathogens and their host cells^27–42^. *T. vaginalis* EVs (TvEVs) can be internalized by both human ectocervical cells and prostate derived cells and have been shown to modulate expression and secretion of the cytokines IL-6 and IL-8^44^. Additionally, pretreatment of human cells with TvEVs can increase cytoadherence of the parasite, an important process in its pathogenesis^44,45^. However, we still know very little about the mechanisms underlying the effects of TvEVs on host cell gene expression and immune signaling pathways. Elucidating the full range of effects of TvEVs on recipient host cells will be important not only to the pathogenesis of *T. vaginalis*, but also to outcomes during *M. hominis* infection as well.

Here we demonstrate that TvEVs suppress interferon-epsilon (IFNε) mediated type I interferon (IFN) responses and are capable of disrupting Signal Transducer and Activator of Transcription 1 and 2 (STAT1/2) nuclear translocation, important host cell transcription factors involved in type I IFN response. Furthermore, we observed that *M. hominis* infection alone or in symbiosis with *T. vaginalis* drives expression of genes involved in the type I IFN response that are not triggered by the parasite itself. Finally, we showed that IFNε is protective against *T. vaginalis*-mediated cytoadherence and cytolysis and enhances neutrophil (PMN) mediated killing of the parasite.

## Results

### Host cells pretreated with *Trichomonas vaginalis* extracellular vesicles are more susceptible to parasite-mediated killing and results in increased parasite burden

TvEVs, produced by the parasite *T. vaginalis,* are known to suppress IL-6 and IL-8 secretion and increase parasite adherence to host cells^44,45^. To determine whether TvEVs also alter parasite-mediated cytolysis of host cells and parasite burden, we modified an automated imaging pipeline to track host cell survival and parasite burden^45^. Human benign prostatic hyperplasia (BPH-1) cells were pretreated in a 96 well plate format with TvEVs or bovine serum albumin (BSA) prior to infection with fluorescently-labeled parasites (Fig. 1a,b). At 24 hours post infection (hpi) entire wells were imaged and host cell survival and parasite burden were quantified (Figs. 1c,d). At 24 hpi we observed an increase in parasite burden when host cells were pretreated with TvEVs compared to BSA control (Fig. 1c). In accordance with increased parasite burden, we also observed a significant decrease in host cell survival when cells were pretreated with TvEVs (Fig. 1d). Together this data indicates that TvEVs increase susceptibility to *T. vaginalis*-mediated cytolysis and increase parasite burden.

**Fig. 1:**
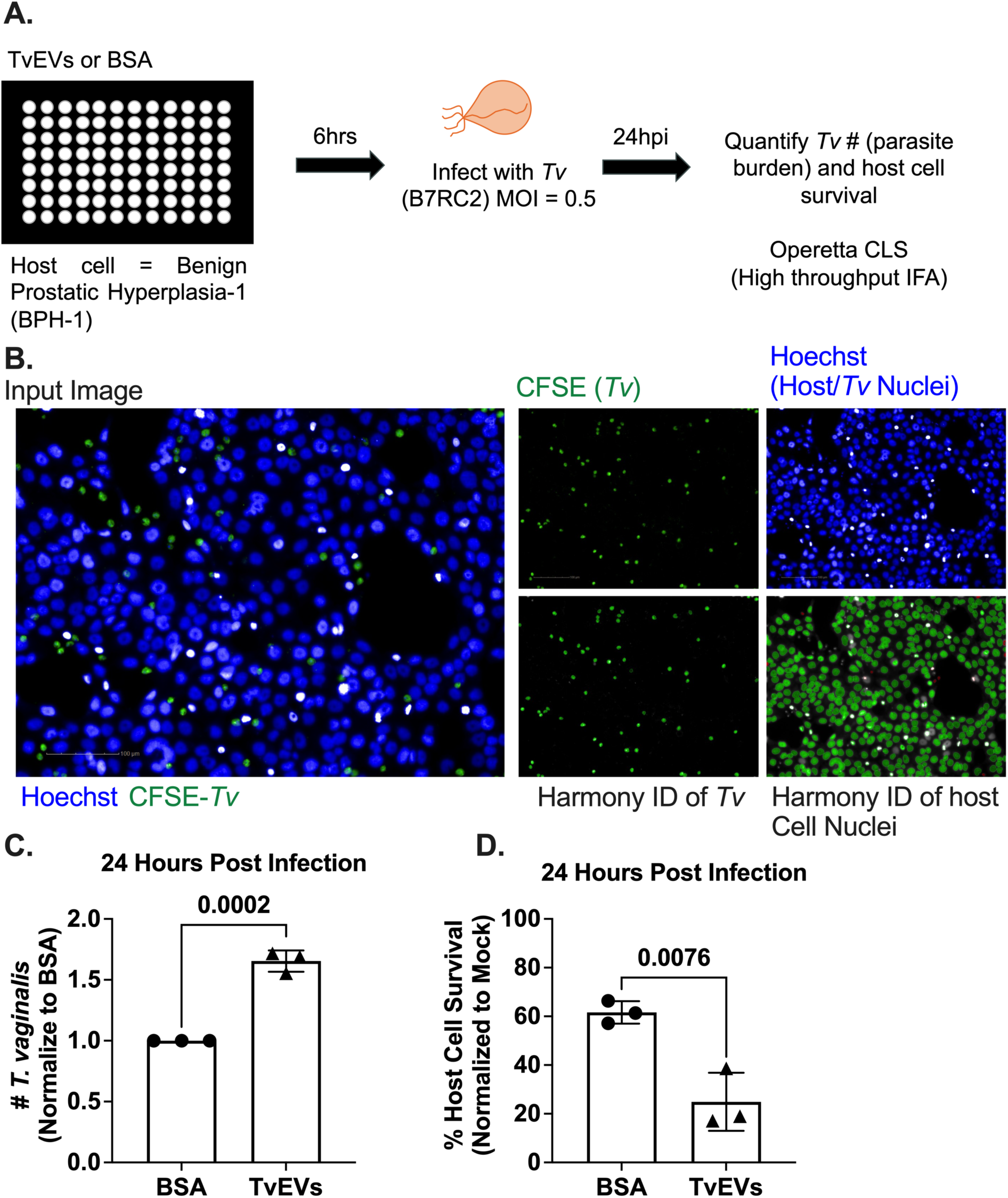
TvEVs increase parasite burden and parasite-mediated host cell cytolysis. **a,** Schematic depicting infection of host cells with *T. vaginalis*. BPH-1 cells were seeded onto 96 well plates to confluency and then treated with 10 µg/mL of TvEVs or BSA for 6 hours prior to infection with CFSE-labeled *T. vaginalis* strain B7RC2 for 24 hours. **b,** Entire wells, treated with either BSA or TvEVs, were imaged on the Operetta^®^ CLS^TM^ platform and analyzed using Harmony™ software. **c,** The number of *T. vaginalis* left in the well after 24 hours was enumerated using CFSE staining to quantify parasites normalized to BSA control. **d,** Percent of host cell survival was quantified by enumerating the number of host cells left in the well after 24 hours compared to uninfected controls. (**c** and **d**) Data is depicted as fold change in adherence compared to BSA control. Bars, mean ± SD. N = 3 wells/experiment, 51 fields of view/well, 3 experiments total. Numbers above bars indicate p-values for unpaired two-tailed Student’s *t*-tests as compared to BSA control.

### Transcriptomic analyses indicate that TvEVs suppress multiple host cell immune pathways, including the type I IFN pathway

We next sought to identify what effect TvEVs have on host cell gene expression as a way of identifying potential mediators of the observed increased susceptibility of host cells to parasite-mediated killing and increased parasite burden (Fig. 1). We treated cells with TvEVs for 6 hours prior to extracting RNA for RNA sequencing (RNA-seq) (Fig. 2a). Sample alignment and quality was assessed using MultiQC (Supplementary Fig. 1)^46^. Transcriptional profiles of the 5 Mock- and 4 TvEV-treated samples were distinguishable by Principle Component Analysis (PCA) (Fig. 2b), with 833 genes (143 upregulated and 690 downregulated) identified as differentially expressed between the two conditions (Fig. 2c,d). Eighty-three percent (690/833) of the differentially expressed genes (DEGs) were downregulated in response to TvEV treatment, indicating a largely suppressive effect on host cell gene expression (Fig. 2c,d). Several of the topmost downregulated DEGs included genes involved in innate immunity including inflammatory cytokines and chemokines (TNFSF14, IL36G, IL36G, CCL5, CCL2, CXCL10, CXCL8) and genes with known antimicrobial properties (NOS2, S100A7, S100A8, S100A9). We next employed Gene Set Enrichment Analysis (GSEA) to identify host cell pathways that were altered^47^. GSEA of the Hallmark dataset indicated that several innate immune signaling pathways (NF-κB signaling via TNFα and interferon alpha and gamma responses) were enriched in mock treated samples compared to TvEV treated samples (Fig. 2e).

**Fig. 2:**
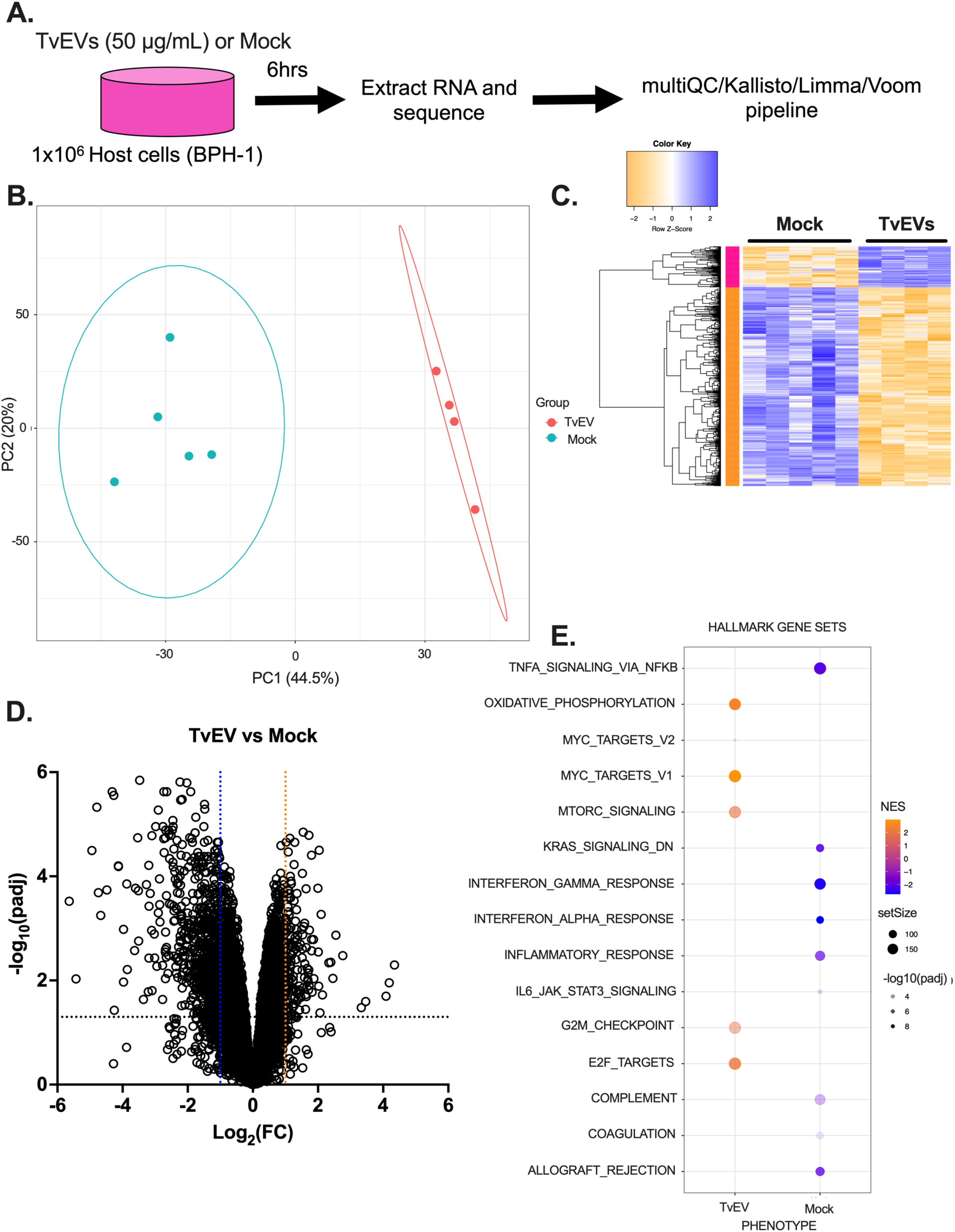
TvEVs down-regulate innate immune genes and pathways. **a,** Schematic depiction of RNA-seq experiment. **b,** Principal Component Analysis (PCA) showing distinct clustering of mock treated BPH-1 cells (Mock; *n* = 5) and TvEV treated BPH-1 cells (TvEV; *n* = 4) from RNA-seq data. **c,** Heatmap of row z-score transformed 832 genes (143 upregulated and 689 downregulated; Log_2_(FC) ≥ 1 or Log_2_(FC) ≤ −1; padj ≤ 0.05) identified as differentially expressed between mock and TvEV treated BPH-1 cells. **d,** Volcano plot depicting differences in gene expression between TvEVs and mock treated BPH-1 cells. The x axis corresponds to the log_2_ fold change in gene expression and the y axis indicates the adjusted p value. Genes with a −log_10_ p value of 1.3 or greater (p value of ≤ 0.05) and a log_2_ fold change −1 ≤ or ≥ 1 were deemed significantly differentially expressed. **e,** Bubble chart showing results of Gene Set Enrichment Analysis (GSEA) for the Hallmarks dataset in TvEV treated (left) and mock treated BPH-1 cells (right). Color of bubble represents normalized enrichment score (gold signifies pathway enriched compared to mock and blue signifies pathway is enriched in mock condition). SetSize indicates number of genes in the indicated pathway. −log_10_(padj) value indicated by intensity of shading.

While previous studies have shown a role for NF-κB signaling in *T. vaginalis* infection^30,48–53^, we were surprised to observe that GSEA predicted that TvEVs suppressed type I and II IFN pathways (Fig. 2e and Extended Data Fig. 1a) as these innate immune pathways are largely characterized for their role in combating intracellular pathogens^54^. However, there is an emerging appreciation for the role of type I IFN responses to certain extracellular pathogens^55^. Closer inspection of our data showed that TvEVs resulted in a significant downregulation of multiple components of the type I IFN signaling pathway (Extended Data Fig. 1b). These included the host cell transcription factors STAT1/2 and Interferon Regulatory Factor 9 (IRF9), which translocate to the host cell nucleus upon binding of the interferon-α/β receptor (IFNAR) with type I IFNs (Extended Data Fig. 1b,c)^54^. The canonical type I IFNs (IFN-α and IFN-β) are secreted in response to stimulation by microbial products and it is possible that TvEVs act as a ligand^29,56,57^. However, transcripts for both IFNA1 and IFNB1 were not detected in our data, instead we observed a significant decrease in the expression of a non-canonical type I IFN, IFNε (Extended Data Fig. 1b,c)^58^. To determine whether TvEVs were capable of disrupting type I IFN signaling, we tested the ability of TvEV pretreatment to block type I IFN responses in cells expressing a luciferase reporter gene downstream of the interferon-stimulated response element (ISRE) sequence^59^. IFNε stimulation resulted in a ∼3.8-fold increase in luciferase activity compared to mock- or TvEV-treated cells (Extended Data Fig. 1d). However, cells pretreated with TvEVs prior to IFNε stimulation resulted in significant diminishment of luciferase activity indicating that TvEVs disrupt type I IFN signaling (Extended Data Fig. 1d).

### Pretreatment of BPH-1 and Ect1 with TvEVs suppress IFNε-mediated nuclear translocation of pSTAT1 and pSTAT2

Our transcriptomic data indicated that STAT1/2 were significantly downregulated by TvEV treatment (Extended Data Fig. 1c). To determine if TvEV treatment also blocked STAT1/2 nuclear translocation, we pretreated BPH-1 with TvEVs prior to stimulating with IFNε and assaying for nuclear translocation of phosphorylated STAT1 and STAT2 (pSTAT1/2). Mock- and TvEV-treated cells displayed little nuclear pSTAT1/2 (Fig. 3a,b). When we stimulated with IFNε we saw a significant increase in nuclear pSTAT1/2 signal confirming that these pathways are active in BPH-1s (Fig. 3c,d). However, pretreatment of host cells with TvEVs prior to IFNε stimulation strongly inhibited pSTAT1/2 nuclear translocation (Fig. 3c,d). These effects were also observed when an ectocervical cell line (Ect1/E6E7) was pretreated with TvEVs prior to IFNε stimulation. In keeping with our BPH-1 data, Ect1s pretreated with TvEVs prior to IFNε stimulation displayed a significant reduction in pSTAT1/2 positive nuclei (Extended Data Fig. 2). Together with our transcriptomic data, these data demonstrate that TvEVs are capable of suppressing type I IFN signaling mediated by IFNε and raises the question of the role of type I IFN responses in *T. vaginalis-*host interactions.

**Fig. 3:**
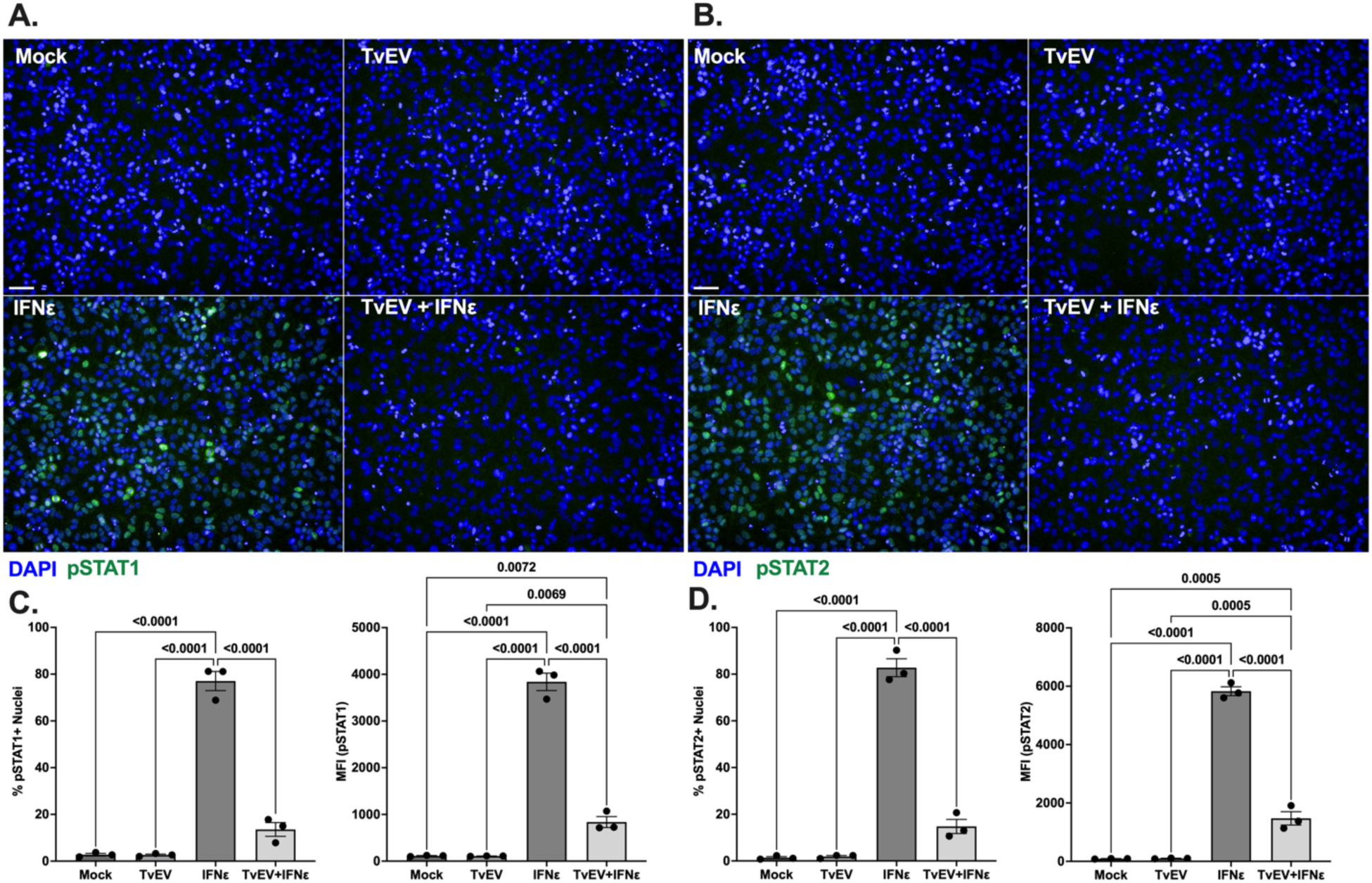
TvEVs block IFNε-mediated nuclear translocation of pSTAT1 and pSTAT2 in prostate cells. **a and b,** pSTAT1/2 immunofluorescence in BPH-1 cells treated with either TvEVs (50 µg/mL), IFNε (500 ng/mL), or TvEVs (50 µg/mL) + IFNε (500 ng/mL). Depicted are DAPI staining BPH-1 and parasite nuclei (Blue) and pSTAT1 or pSTAT2 (Green). **c,** *Left,* quantification of percent of cells that at positive for nuclear pSTAT1. *Right,* Mean Fluorescent intensity (MFI) of nuclear pSTAT1. **d,** *Left*, quantification of percent of cells that at positive for nuclear pSTAT1. *Right*, Mean Fluorescent intensity (MFI) of nuclear pSTAT2. (**c** and **d**) Bars, mean ± SD. N = 3 wells/experiment, 9 fields of view/well, 3 experiments total. Numbers above bars indicate p-values for one-way ANOVA, Dunnett’s multiple comparisons test compared to vehicle control.

### *T. vaginalis* and its bacterial symbiont, *M. hominis,* elicit distinct and overlapping responses in host gene expression during infection

While there is an emerging appreciation for the role of type I IFN responses to extracellular pathogens, the type I IFN response has historically been studied for its role in combating intracellular pathogens^54,55^. Interestingly, some strains of *T. vaginalis* are known to harbor up to four different double-stranded RNA viruses, *Trichomonas vaginalis* virus 1-4 (TVV1-4)^19,60^, as well as a bacterial symbiont *M. hominis,* which is an opportunistic intracellular pathogen of the female reproductive tract (FRT)^19,61^. The possibility that TVV might be contributing to the type I IFN responses we observed during *T. vaginalis* infection can be eliminated as the parasite strain used here (B7RC2) is free of TVV1-4^45^. However, *T. vaginalis* strain B7RC2 does harbor the *M. hominis* symbiont (Supplementary Fig. 2).

To determine how *T. vaginalis*, *M. hominis*, and *T. vaginalis* in symbiosis with *M. hominis* affected host cell gene expression we infected BPH-1 with either *T. vaginalis* that had been cleared of *M. hominis* (Tv), *M. hominis* isolated from *T. vaginalis* (Mh), or *T. vaginalis* in symbiosis with *M. hominis* (TvMh) and then extracted RNA from host cells and performed RNA-seq. Tv samples were confirmed clear of *M. hominis* by PCR (Supplementary Fig. 2). Sample alignment and quality was assessed using MultiQC (Supplementary Fig. 3)^46^. Transcriptional profiles of 5 Mock, 5 Tv infected, 5 Mh infected, and 5 TvMh infected samples were distinguishable by PCA with the TvMh infected samples clustering between the Tv and Mh samples (Extended Data Fig. 3). DEGs fell into distinct modules of expression corresponding to infection with either Tv (green module) or Mh (pink module) or TvMh (Extended Data Fig. 4a). We identified 2294 DEGs (1134 upregulated and 1160 downregulated) in Tv infected cells, 1314 DEGs (637 upregulated and 677 downregulated) in Mh infected cells, and 2886 DEGS (1438 upregulated and 1448 downregulated) in TvMh infected cells, compared to mock controls (Extended Data Fig. 4b-d). Of the 1619 upregulated DEGs, 45.4% (735/1619; **I** and **IV** of Venn diagram) were specific to Tv infected samples, 14.7% (238/1619; **III** and **VI** of Venn diagram) to Mh infected samples, and 15.3% (247/1619; **VII** of Venn diagram) to TvMh infected samples (Extended Data Fig. 4e). Of the 1806 downregulated DEGs, 39.9% (721/1806; **I** and **IV** of Venn diagram) were specific to Tv infected samples, 13.2% (238/1806; **III** and **VI** of Venn diagram) to Mh infected samples and 22.6% (408/1806; **VII** of Venn diagram) to TvMh infected samples (Extended Data Fig. 4f). Of the 1619 upregulated and 1806 downregulated DEGs 24.3% (394/1619) and 22.4% (405/1806), respectively, were common to all infections (Extended Data Fig. 4e, f). Together these data indicate that while *T. vaginalis* and *M. hominis* elicit distinct transcriptional responses from host cells during infection there is also a large overlap on host cell gene expression.

### *T. vaginalis* association with *M. hominis* drives enrichment of the type I IFN pathway

GSEA of Tv infected cells revealed that, in contrast to the response to TvEVs (Fig. 2), there was an enrichment of several innate immune signaling pathways (NF-κB signaling via TNFα, inflammatory response, complement, and interferon gamma response) (Fig. 4a Extended Data Fig. 5) suggesting that TvEVs may be released by the parasite to suppress innate immune responses induced by infection. One exception to this trend was the negative normalized enrichment score for the interferon alpha response pathway upon treatment with TvEVs. In contrast, GSEA of Tv infected host cells did not have a significant enrichment in the interferon alpha response (Fig. 4a and Extended Data Fig. 5). However, Mh infected and TvMh infected cells were significantly enriched in the interferon alpha response pathway compared to mock infected cells (Fig. 4a and Extended Data Fig. 5).

**Fig. 4:**
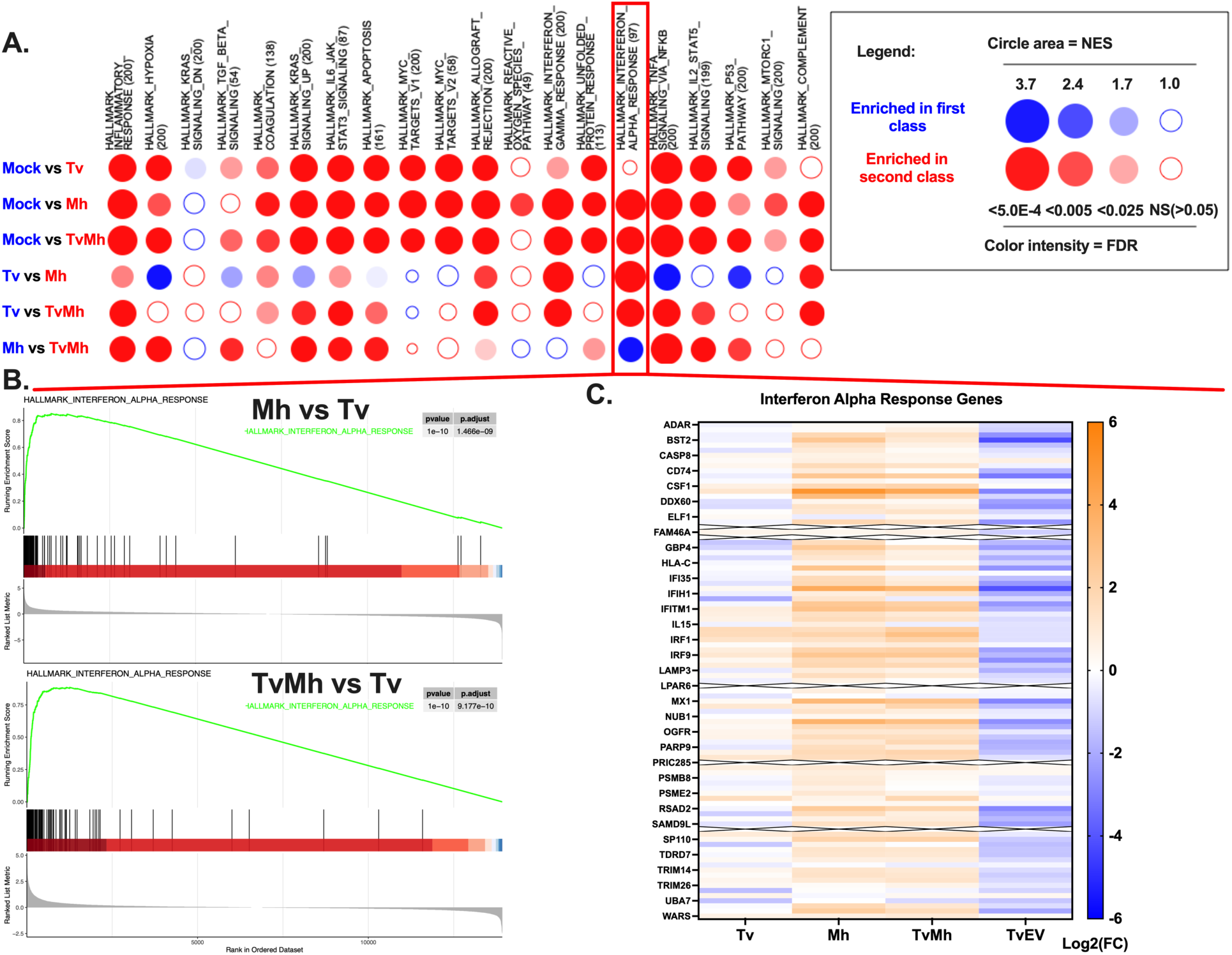
GSEA reveals that *T. vaginalis* association with *M. hominis* drives enrichment of the IFNα response pathway. **a,** BubbleMAP showing results of Gene Set Enrichment Analysis (GSEA) for immune pathways in Hallmarks dataset for BPH-1 cells infected with Tv, Mh, or TvMh. Color and area of bubble represents normalized enrichment score (NES). Blue signifies pathway enriched in first class and Red signifies enrichment in second class. Intensity of color indicates false discovery rate (FDR). **b,** *Top*, GSEA enrichment plot showing Hallmark IFN alpha response of Mh infected BPH-1 cells compared to Tv infected BPH-1 cells. *Bottom,* GSEA enrichment plot showing Hallmark IFN alpha response of TvMh infected BPH-1 cells compared to Tv infected BPH-1 cells**. c,** Heatmap of Log_2_(FC) of individual genes in the interferon alpha response pathway Hallmarks data set for samples normalized to Mock control. Rows with an X through them represent genes with transcripts below limit of detection in the RNA-seq data.

We reanalyzed our transcriptomic data to focus on differences in gene expression between Mh and TvMh infected cells compared to Tv infected cells. We identified 557 DEGs (213 upregulated and 344 downregulated) in Mh infected cells compared to Tv infected cells and 171 DEGs (170 upregulated and 1 downregulated) in TvMh infected cells compared to Tv infected cells (Extended Fig. 6). GSEA revealed that Mh and TvMh infected cells were enriched in the interferon alpha and gamma responses compared to Tv infected cells (Fig. 4a,b and Extended Data Fig. 5). Additionally, analysis of the individual genes that comprise the interferon alpha response pathway in the Hallmarks dataset showed a general upregulation of gene expression in Mh and TvMh infected cells compared to mock infected cells, while Tv infection or TvEV treatment showed a general suppression of genes in the interferon alpha response pathway (Fig. 4c). Together our transcriptomic datasets indicate that *M. hominis* is likely the main driver of type I IFN responses during *T. vaginalis* infection and that these responses may be dampened by secretion of TvEVs.

### *T. vaginalis* cytoadherence and host cell cytolysis is inhibited by IFNε-mediated type I IFN responses

Our data show that TvEVs are capable of dampening type I IFN responses, and while we were unable to detect transcripts for IFN-α and IFN-β in our data, we did detect transcripts for IFNε, and observed a significant decrease in IFNε gene expression in TvEV treated cells compared to mock controls (Extended Data Fig. 1b,c). IFNε has previously been shown to play an important role in protection of the FRT from both bacterial and viral STIs and is expressed by epithelial cells of the FRT and prostate^58,62–68^. Interestingly, IFNε, unlike other interferons, is not regulated by pattern recognition receptor (PRR) stimulation, but rather is hormonally regulated which may explain the high basal level of expression in the BPH-1 cell compared to IFN-α and IFN-β^58^. Epithelial cells at mucosal surfaces such as the FRT and male urogenital tract (MUT) are often the first cells to come into contact with invading pathogens and it is thought that tissue specific expression of cytokines like IFNε may be involved in priming these cells for protection against pathogens^69^.

To determine whether IFNε is protective against *T. vaginalis,* we stimulated BPH-1s with IFNε prior to infection with parasites (Fig. 5a,b). At 2 hpi no differences in host cell survival or parasite burden was observed between stimulated and mock-treated cells (Fig. 5c,d). However, at 24 hpi we observed an increase in host cell survival and a decrease in parasite burden for cells pretreated with IFNε (Fig. 5c,d). We also found that the protective effects of IFNε was inhibited by pretreatment with either TvEVs, an IFNAR1 blocking antibody, or a JAK inhibitor prior to IFNε stimulation and subsequent *T. vaginalis* infection, indicating that TvEVs or inhibition of proteins involved in the type I IFN pathway disrupts IFNε-mediated protection (Extended Data Fig. 7).

**Fig. 5:**
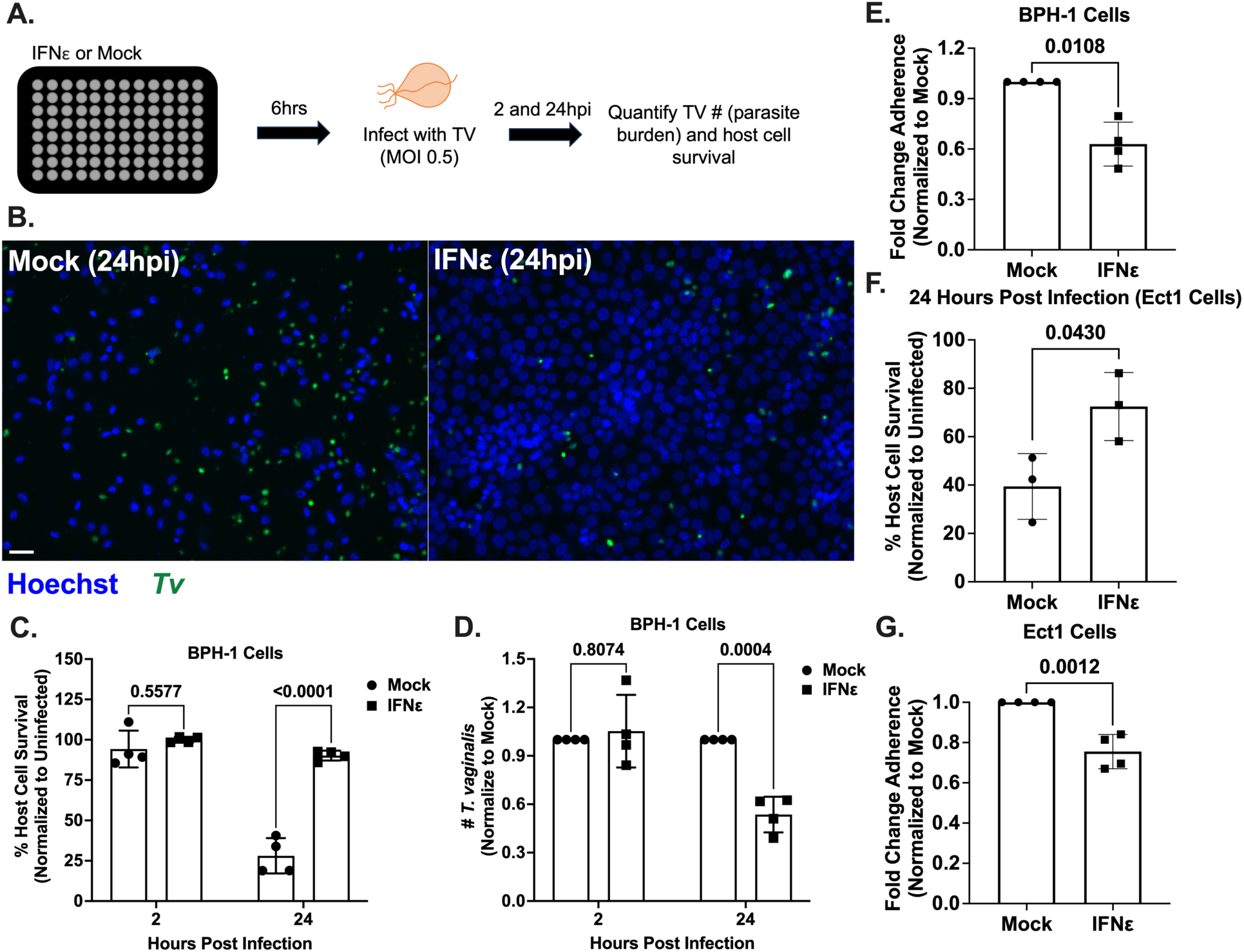
IFNε is protective against *T. vaginalis* cytoadherence and cytolysis. **a,** *Top,* Schematic depicting infection of host cells with *T. vaginalis*. BPH-1 cells were seeded onto 96 well plates to confluency and then treated with 500 ng/mL of IFNε or mock treated for 6 hours prior to infection with CFSE-labeled *T. vaginalis* strain B7RC2 for 2 and 24 hours. **b,** Representative fluorescence image depicting BPH-1 cells mock or IFNε (500 ng/mL) treated and then infected with *T. vaginalis* at 2 and 24 hours post infection. Hoechst staining for BPH-1 and parasite nuclei (Blue) and CFSE-labeled *T. vaginalis* (Green). **c,** Percent of host cell survival was quantified by enumerating the number of host cells left in the well after 2 and 24 hours compared to uninfected controls. **d,** The number of *T. vaginalis* left in the well after 2 and 24 hours was enumerated using CFSE staining to quantify parasites normalized to Mock treated control. **e,** Quantification of parasite adherence to BPH-1. Data is depicted as fold change in adherence compared to mock treated control. **f,** Percent of host cell survival was quantified by enumerating the number of host cells left in the well after 24 hours compared to uninfected controls. **g,** Quantification of parasite adherence to Ect1. Data is depicted as fold change in adherence compared to mock treated control. (**c** and **d**) Bars, mean ± SD. N = 3 wells/experiment, 51 fields of view/well, 4 experiments total. Numbers above bars indicate p-values for two-way ANOVA, Dunnett’s multiple comparisons test compared to mock control. (**e**) Bars, mean ± SD. N = 3 wells/experiment, 4 experiments total. Numbers above bars indicate p-values for Welch’s t-test compared to mock control. (**f** and **g**) Bars, mean ± SD. N = 3 wells/experiment, 3 experiments total. Numbers above bars indicate p-values for Welch’s t-test compared to mock control.

*T. vaginalis-*mediated cytolysis has been shown to be contact dependent^70^. Thus, we decided to test if the protective effect of IFNε might be mediated in part by decreased parasite adherence to host cells. When BPH-1 was stimulated with IFNε prior to performing an adherence assay, we observed a ∼40% reduction in the number of parasites adhering to IFNε stimulated cells compared to mock controls (Fig. 5e). We then determined whether IFNε would also be protective against parasite-mediated cytolysis and adherence in Ect1s. Consistent with our BPH-1 data (Fig. 5c), IFNε stimulation resulted in a significant increase in Ect1 survival (Fig. 5f) and a decrease in parasite adherence (Fig. 5g). These data demonstrate that IFNε is protective against parasite cytoadherence and cytolysis (Fig. 5) and that the protective effects of IFNε are mediated through signaling of the type I IFN pathway (Extended Data Fig. 7) and can be disrupted by TvEVs (Extended Data Fig. 7).

### IFNε enhances PMN-mediated killing of *T. vaginalis*

Studying the *in vivo* immune response of the FRT to *T. vaginalis* has been severely limited due to the lack of a robust female mouse model of infection. However, clinical observation and *in vitro* work has identified neutrophils (PMNs) as important immune cells in the response to *T. vaginalis* infection^71–73^. Clinical presentation of trichomoniasis is associated with the influx of PMNs to the vaginal mucosa and PMNs have been shown to kill *T. vaginalis* by trogocytosis^72,73^. Release of neutrophil extracellular traps (NETs) has also been reported to play a role in protection against *T. vaginalis*^74^. This led us to ask whether IFNε stimulation might play a role in PMN-mediated killing of *T. vaginalis*. To this end, we pretreated human PMNs with IFNε prior to co-incubation with *T. vaginalis* and assayed for parasite death (Supplementary Fig. 4). We observed significantly increased parasite death with IFNε stimulation of PMNs, demonstrating that IFNε can increase the parasite killing capacity of PMNs (Fig. 6). These data demonstrate that IFNε stimulation increases *T. vaginalis* killing by PMNs involved in parasite clearance during infection.

**Fig. 6:**
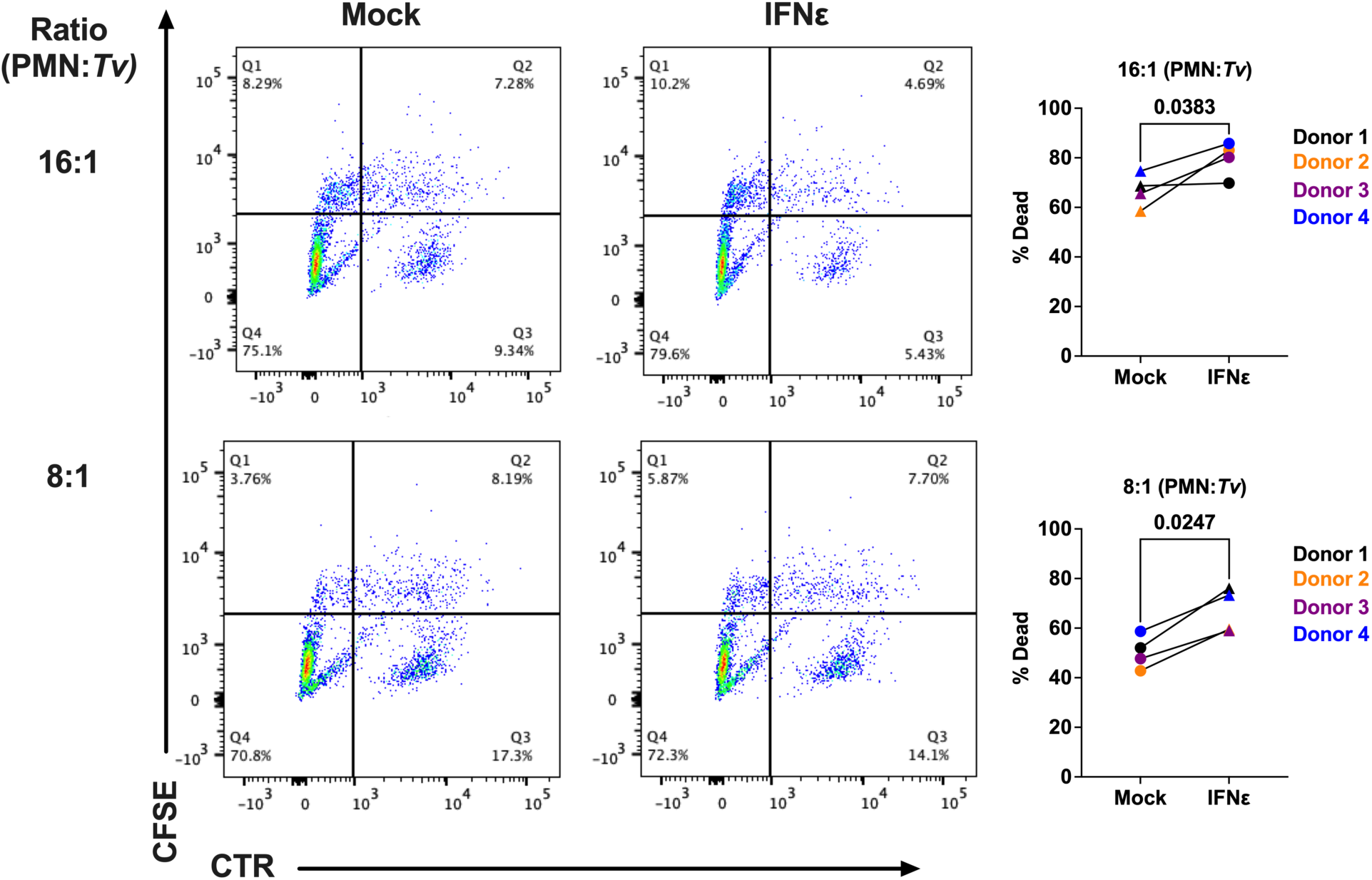
IFNε increases PMN-mediated killing of *T. vaginalis.* Scatter plot for 1 individual donor showing remaining live population Cell Tracker Red (CTR) labeled *T. vaginalis* after a 4 hour incubation with CFSE-labeled PMNs mock treated or pre-stimulated with IFNε (1 µg/mL) at 16:1 and 8:1 (PMN:*Tv*) ratio. Percent death of *T*. *vaginalis* was calculated as number of *T. vaginalis* (Q3) left after co-culture with PMNs divided by the number of *T. vaginalis* left after 4 hours with no PMNs multiplied by 100. Each dot represents 1 unique donor (4 donors total) with lines connecting the same donor per condition. Numbers above bars indicate p-values for unpaired T-test compared to mock control

## Discussion

Extracellular vesicles (EVs) secreted by many microorganisms have been shown to modulate host cell immune responses during infection^75,76^. Here we demonstrate a novel connection between type I IFN responses mediated by IFNε and TvEVs. Specifically, we showed that TvEVs released by the parasite suppress innate immune signaling pathways and can disrupt type I IFN responses resulting in an increase in host cell death. Furthermore, transcriptional analysis of host cells infected with *T. vaginalis, M. hominis*, or *T. vaginalis* in symbiosis *M. hominis* revealed that type I IFN responses are largely elicited in response to *M. hominis*, but not *T. vaginalis*. Finally, we demonstrated that IFNε is protective against parasite-mediated cytolysis and enhances the ability of PMNs to kill *T. vaginalis*.

TvEVs are internalized by human cells resulting in increased parasite adherence to host cells and altered cytokine secretion^37,44^. Here we show that pretreatment of human cells with TvEVs results in an increase in both host cell cytolysis and parasite burden. These observations led us to interrogate the specific effects of TvEVs on host cell gene expression, revealing that TvEVs largely suppressed host cell gene expression and downregulated many immune related genes and pathways. Multiple genes involved in the type I IFN pathway were significantly downregulated by TvEV treatment. These effects translated into an ability to inhibit IFNε-mediated activation of the type I IFN response and blocked nuclear translocation of pSTAT1/2. These findings suggest that TvEV cargo (i.e., parasite proteins, small RNAs, or other small molecules) inhibit activation of the type I IFN response. Proteomic analyses have identified several TvEV proteins that could mediate downstream host cell signaling (i.e. kinases, phosphatases, and phospho-binding proteins)^44,57^. Alternatively, the immunosuppressive effects of TvEVs may be the result of small RNAs present in TvEVs, as described for EVs from helminthic parasites known to carry micro-RNAs that modulate the expression of host cells genes involved in the immune response^31,34,42,77–79^. We can eliminate the possibility that TVV RNAs are involved, as previously reported for TvEV-mediated downregulation of IL-6 and IL-8, as the *T. vaginalis* strain used for these studies does not harbor TVV^45^. Future studies will be necessary to identify the exact TvEV cargo that mediate the suppressive effects of TvEVs on the type I IFN response.

Type I IFN responses have primarily been implicated in combating intracellular pathogens and not extracellular parasites such as *T. vaginalis*^54^. Notably, a high percentage of *T. vaginalis* strains harbor a bacterial symbiont, *M. hominis*^19^. It is unknown whether the symbiont alters *T. vaginalis* clinical outcomes. In addition to showing that TvEVs suppress type I IFN responses, we have demonstrated that the enrichment of type I IFN response pathway is only seen in host cells infected with *T. vaginalis* harboring *M. hominis* or during *M. hominis* infection alone, but not in infections with the same *T. vaginalis* strain cleared of *M. hominis*. These findings indicate that *M. hominis* activates the type I IFN response observed in infections when the parasite harbors this symbiont. Such a possible negative consequence of this symbiosis is likely balanced by the benefits the bacterium confers to *T. vaginalis* by increasing parasite growth, hemolytic activity, and adherence to host cells^18,27,80–82^. We propose that to counter the negative consequence of the symbiont activating the type I IFN pathway, the parasite secretes TvEVs that suppress this innate immune response. Future work elucidating and defining the immune response to *T. vaginalis* and *M. hominis* will be important in revealing how symbiosis alters the pathogenesis of both microorganisms.

In addition to demonstrating that TvEVs can suppress type I IFN responses, we have shown that activation of the type I IFN response by IFNε prior to infection is protective against *T. vaginalis-* mediated cytolysis of host cells. This is particularly interesting as IFNε plays an important role in protection of the FRT from both bacterial and viral STIs and is expressed by epithelial cells of the FRT and the MUT. IFNε has a protective role against herpes simplex virus 2, zika virus, HIV, and *Chlamydia muridarum*, and has recently been implicated in restriction of ovarian cancer^58,62,64,68,83^. How IFNε mediates protection of human cells against *T. vaginalis-*mediated adherence and cytolysis remains to be elucidated. It may be that IFNε stimulation alters critical cell surface components of host cells required for parasite adherence. Alternatively, IFNε stimulation may result in the release of molecules with antiparasitic activity.

It is well established that neutrophils/PMNs are the most abundant immune cells present at the site of *T. vaginalis* infection and previous work has demonstrated that PMNs kill *T. vaginalis* by early stage trogocytosis and late stage NETosis^71–74,84^. For other pathogens IFN*α* and IFN*β* are known to enhance recruitment of PMN to site of infection, regulate neutrophil function, delay apoptosis, and trigger the release of NETs^85^. However, a possible role for IFNε in PMN-mediated killing of pathogens has not been reported. Here we show that pretreatment of PMNs with IFNε results in an increased ability of PMNs to kill *T. vaginalis*, suggesting an activating role for IFNε in PMNs. How IFNε stimulation alters PMNs and their ability to kill *T. vaginalis* remain to be determined.

Our observation that IFNε protects host cells from *T. vaginalis* infection also sheds light on epidemiological studies showing an unusually high prevalence of *T. vaginalis* infection in peri/post- menopausal aged women. In contrast to other STIs, the detection rate of *T. vaginalis* does not decrease with age and reaches maximum rates in women 48-51 years old, raising the possibility that physiological and immunological changes due to menopause may make the FRT more conducive to infection^16,86^. Interestingly, IFNε, unlike other IFNs, is hormonally regulated by estrogen and progesterone^65^. In mice, IFNε levels during estrus are 30-fold higher than in diestrus and in humans IFNε expression is 10-fold higher during the proliferative phase of the menstrual cycle (late follicular) compared to the secretory phase (luteal)^58^. Notably, IFNε levels are ∼100-fold decreased in epithelial cells of the FRT from postmenopausal women raising the possibility that loss of IFNε expression may play a role in the increased association of *T. vaginalis* positivity in aging women^58,63^.

The results presented here have demonstrated that infection with *T. vaginalis* in symbiosis with *M. hominis* increases expression of genes in the type I IFN response and show that IFNε, which is greatly diminished in peri/post-menopausal women, is protective against *T. vaginalis* infection. Additional distinct and overlapping immune pathways were found to be stimulated by the parasite dependent on the presence and absent of the symbiont. We also identified TvEVs as a means used by the parasite to suppress innate immune responses and enhance host cell killing. Elucidating the mechanisms that drive these dynamic interactions between this human parasite, its symbiotic bacterium, and the host cell will lead the way towards a fuller understanding of this prevalent STI.

## Methods

### Parasite Maintenance

*T. vaginalis* strain B7RC2^70^ was cultured in Diamond’s modified Trypticase-yeast extract-maltose (TYM) medium supplemented with 10% horse serum (Sigma-Aldrich), 10 U/ml penicillin and 10 μg/ml streptomycin (Gibco), 180 μM ferrous ammonium sulfate, and 2 μM sulfosalicylic acid. To generate *M*. *hominis* free strains, *Tv* was grown in the presence of 50 μg/ml chloramphenicol and 5 μg/ml tetracycline (Sigma-Aldrich), supplemented daily for at least 5 days^87^. *M*. *hominis* clearance was confirmed by PCR as described below prior to use in experiments^87,88^. Parasites were grown at 37°C and passaged daily for no more than 2 weeks at a time.

### Human Cell Culture

Human benign prostate hyperplasia 1 (BPH-1) epithelial cells^89^ were cultured at 37°C with 5% CO_2_ in RPMI 1640, L-glutamine, and HEPES media (Gibco) supplemented with 10 U/ml penicillin, 10 μg/ml streptomycin, and 10% fetal bovine serum (FBS; Gibco) as previously described^44^. The human ectocervical cell line Ect1 E6/E7^90^ (CRL 2614) was grown in keratinocyte-SFM (KSFM) (Invitrogen) supplemented with 10 U/ml penicillin, 10 μg/ml streptomycin, and 0.4M Ca^2+^ as described previously^44^.

### Extracellular Vesicle Isolation

*T. vaginalis* was grown to confluency (1×10^6^ cells/mL) in 50 mLs of complete Diamond’s media. Parasites were spun down at 3200 rpm for 10 minutes and washed with 50 mLs serum-free Diamond’s media twice. Cells were transferred to 50 mLs serum-free Diamond’s media and incubated at 37°C for 4 hours. Cells were spun down at 3000 rpm at 4°C for 20 minutes to remove cell debris. The supernatant was kept and mixed 2:1 with Total Exosome Isolation Reagent (Invitrogen #4478359). The supernatant mixture was incubated overnight at 4°C. Precipitated TvEVs were recovered by standard centrifugation at 10,000xg for 60 minutes at 4°C. The exosome pellet was resuspended in sterile 1x PBS plus 1:100 Halt Protease Inhibitor Cocktail (Thermo Scientific #78428) and stored at −80°C. The concentration of TvEV protein was determined using Pierce BCA Kit (Thermo Scientific #23225). To generate the mock control used in our transcriptomic analyses we used 50 mLs serum-free Diamond’s alone (no parasites) subjected to the isolation procedure described above^45^.

### Host Cell Survival / Parasite Burden Assay

Host cell survival and parasite burden of BPH-1 and Ect1 cells infected with *T. vaginalis* was performed using a modified protocol for measuring parasite adherence in a 96-well plate format on the Operetta^®^ CLS^TM^ platform^45^. CFSE-labeled parasites were added to confluent monolayers of BPH-1 or Ect1 cells at an MOI of 0.5 at the indicated time points. At the indicated time points cells and parasites were fixed in 4% formaldehyde in 1x PBS for 20 minutes at room temperature. Host cells and parasites were stained with Hoechst (1:1000) (Biotium; Cat. #40044) for 15 minutes at room temperature. Plates and entire wells were imaged using the Operetta^®^ CLS^TM^ platform, high throughput microplate imaging system and analyzed using Harmony™ software. CFSE positive parasites were identified using the Harmony™ identify cells tool. Fluorescent intensity cutoffs were set at 1,000 and length and width cutoffs were set at 5 μms and 2 μms respectively to filter out any small auto-fluorescent debris. Host cell nuclei were identified by using the Harmony™ identify nucleus tool. Parasite burden was determined as the number of parasites in the well relative to BSA or mock controls. Host cell survival was quantified by enumerating the number of host cells left in the well after 24 hours compared to uninfected controls. For anti-IFNAR blocking assays cells were preincubated with 1 µg/mL of Anti-interferon alpha/beta receptor 1 antibody (Abcam; Cat # ab124764) or IgG control for 1 hour prior to infection with parasites. For JAK inhibitor assays cells were pretreated with 10 µM of Ruxolitinib (Thermo Scientific; Cat # AC469381000) or mock treated (DMSO control) for 1 hour prior to infection with parasites.

### Transcriptomic Analysis

For TvEV transcriptomic analysis BPH-1 cells (1×10^6^ cells per condition per replicate) were treated with 50 µg/mL of TvEVs or mock treated (see EV Isolation methods section) for 6 hours in serum free RPMI prior to RNA isolation. For transcriptomic analysis of infected BPH-1, 1×10^6^ cells per condition per replicate were infected (MOI = 1) with *T. vaginalis* that had been cleared of *M. hominis* (Tv) using previously established protocols, *M. hominis* isolated from culture with *T. vaginalis* (Mh), *T. vaginalis* in symbiosis with *M. hominis* (TvMh), or mock infected for 6 hours in serum-free RPMI media at 37°C prior to total RNA isolation. Total RNA was isolated using the Direct-zol^TM^ RNA Miniprep Kit (Zymo Research; Cat. #R2071) according to the manufacturer’s instructions. Total RNA was sent to Novogene for RNA quality control (QC) analysis, library prep and sequencing. RNA QC was analyzed, and quantification of RNA preparations and libraries were carried out using an Agilent 5400. Samples were sequenced on an Illumina NovoSeq 6000 to produce 150–base pair paired end reads with a mean sequencing depth of 20 million reads per sample. After read mapping with Kallisto^91^, version 0.46.2, TxImport^92^ was used to read Kallisto outputs into the R environment. Annotation data from Ensembl (https://ftp.ensembl.org/pub/release-110/gtf/homo_sapiens/) was used to summarize data from transcript-level to gene-level. All subsequent analyses were carried out using the statistical computing environment R version 4.3.0 in RStudio version 1.1.456, and Bioconductor version 3.8. Briefly, transcript quantification data were summarized to genes using the tximport package and normalized using the trimmed mean of M values (TMM) method in edgeR^93^. Genes with <1 CPM in *n* + 1 of the samples, where *n* is the size of the smallest group of replicates, were filtered out. Normalized filtered data were variance-stabilized using the voom function in limma^94^, and differentially expressed genes were identified with linear modeling using limma (FDR ≤ 0.05; absolute Log_2_FC ≥ 1) after correcting for multiple testing using Benjamini-Hochberg. GSEA analysis was performed using the BubbleGUM package as previously described^95^.

### PCR Verification of *M. hominis*

To assay for the presence of *M. hominis* in our *T. vaginalis* strains total genomic DNA was isolated from 15 mL overnight cultures as previously described. Genomic DNA was then used in a PCR reaction with *M*. *hominis* specific primers: 5’-CAATGGCTAATGCCGGATACGC-3’ and 5’- GGTACCGTCAGTCTGCAAT-3’ as previously described^87,88^.

### M. hominis isolation from T. vaginalis culture

*M. hominis* was isolated from culture with *T. vaginalis* for use in RNA-seq experiments as follows. 50 mLs of overnight *T. vaginalis* culture was centrifuged at 3200 rpm for 10 minutes to pellet *T. vaginalis,* while extracellular *M. hominis* remained in the supernatant. Supernatant was gently transferred to a new 50 mL conical tube and subjected to repeated centrifugation 2 times. After the third centrifugation 10 µLs of supernatant was visualized under a light microscope to ensure no parasites remained.

### Type I IFN-Activated Luciferase Reporter Assay

HeLa cells stably expressing 11×-ISRE-Gaussia Luciferase reporters were kindly gifted to us by Dr. David Sibley^59^. Luciferase reporter cells were treated with 10 µg/mL of B7RC2 TvEVs for 6 hours and then stimulated with IFNε (500 ng/mL) for another 6 hours. The cells were then washed and assayed for expression of Gaussian Luciferase in cell lysates using a BioLux Gaussia Luciferase Assay Kit (New England BioLabs) as per the manufacturer’s instructions.

### pSTAT1 and pSTAT2 Nuclear Translocation by Immunofluorescence

BPH-1 or Ect1 cells were treated with either TvEVs (50 µg/mL) or mock treated for 6 hours and then stimulated with IFNε (500 ng/mL) for 1 hour. The cells were fixed with 100% methanol for 15 min and blocked with 5% w/v BSA for 1 hour. Fixed cells were stained with 1:200 rabbit anti-phospho-STAT1 (Cell Signaling Technologies) or 1:200 rabbit anti-phospho-STAT2 (Cell Signaling Technologies) overnight at 4 °C, followed by 1:1000 goat anti-rabbit Alexa 488 (Thermofisher), or 1:1000 goat anti-mouse Alexa 568 (Thermofisher), and 5 μg/mL DAPI for 1 h. Images were acquired at 20x on an Operetta^®^ CLS^TM^ platform, high throughput microplate imaging system and analyzed using Harmony™ software. Briefly, nuclei in each sample were identified as primary objects using DAPI staining and the find nucleus tool, mean pSTAT1/2 intensity in identified nuclei was measured as the nuclear pSTAT1/2 level. Nuclei were deemed positive if they had a pSTAT1/2 signal of MFI ≥ 1000.

### Adherence Assay

Adherence of *T. vaginalis* to BPH-1 cells was performed as described previously in a 96-well plate format on the Operetta^®^ CLS^TM^ platform^45^. 1 × 10^6^ CFSE-labeled *T. vaginalis* parasites were incubated with the indicated amount of TvEVs for 30 minutes and then 5 × 10^4^ parasites were added to confluent monolayers of BPH-1s for 1 hour. Unattached parasites were washed off and cells were fixed in 4% formaldehyde in 1x PBS for 20 minutes at room temperature. Plates were imaged using the Operetta^®^ CLS^TM^ platform, high throughput microplate imaging system and analyzed using Harmony™ software. CFSE positive parasites were identified using the Harmony™ identify cells tool. Fluorescent intensity cutoffs were set at 1,000 and length and width cutoffs were set at 5 μm and 2 μm respectively to filter out small auto-fluorescent debris. The % attachment was determined as the number of parasites in the well divided by input (5 × 10^4^) multiplied by 100.

### PMN Isolation

Polymorphonuclear neutrophils (PMNs) from four random, de-identified donors from the UCLA Health Blood and Platelet Center were extracted as follows^84^. Whole blood was processed using a Leukocyte Reduction System (LRS) filter to remove platelets. The remaining blood was diluted 1:5 with DPBS without Ca^2^+/Mg (Thermo Fisher Scientific, Carlsbad, CA) placed on top of Ficoll-Paque PLUS (Cytiva). PMNs (pellet) were separated from peripheral blood mononuclear cells (PBMC) (supernatant) by density gradient isolation via centrifugation at 150xg for 20 minutes at room temperature with no breaking. The pellet was then resuspended in HBSS (Hanks’ Balanced Salt Solution without Ca^2^+/Mg, Gibco) and an equal volume of 3% Dextran (Pharmacosmos, Denmark) in 0.9% NaCl and incubated until two distinct levels form, 30-60 minutes. PMNs in the top layer were removed from erythrocytes in the bottom layer and centrifuged at 250xg for 10 minutes at room temperature. The PMN pellet was resuspended in 20 mLs ACK lysing buffer for 5 minutes (Invitrogen; Thermo Fisher Scientific, Carlsbad, CA) to lyse any remaining erythrocytes. Samples were subsequently centrifuged at 300xg for 5 minutes and the PMN pellet was washed in 10 mLs DPBS and cells were immediately counted before fluorescent labelling with CFSE.

### PMN Parasite Killing Assay

PMN-mediated parasite killing was performed following modified methods described previously^84^. Briefly, CFSE labelled PMNs were pre-treated with 1 µg/mL of IFNε for 4 hours or mock treated prior to adding Cell Tracker Red (CTR) labelled *T. vaginalis* at ratios of 16:1 and 8:1 (PMN:*Tv*), keeping the *T. vaginalis* count constant at 15,000 parasites per well. The co-cultures were left for an additional 4 hours in a 96 well format in RPMI supplemented with 10% hABS at 37° C in 5% CO_2_. After 4 hours cells were fixed with 4% formaldehyde and parasite survival was measured by counting the live CTR labelled parasites via Flow Cytometry.

### Statistical Analysis

Graphs were generated and statistical analyses were performed using Prism 10.0.2 software. All experiments were performed at least three independent times, with at least 3 technical replicates per experiment, and statistical analyses were conducted on the composite data unless reported otherwise.

## Data Availability

Raw sequence data are available on the Gene Expression Omnibus (GEO; GSE278319 and GSE278360). All other data are included in the manuscript and/or supplemental information.

## Acknowledgements

We want to thank our colleagues in the Johnson lab and at UCLA for their helpful discussions and critiques. This research was supported by NIH grants R01AI103182 and R01AI148475 (to P.J.J.). J.A.K. received support from NIH Ruth L. Kirschstein National Research Service Award T32AI007323 and Ruth L. Kirschstein National Research Service Award Individual Fellowship F32AI186416. J.A. and G.E. received support from National Summer Undergraduate Research Project, NSF Grant #: DBI-2149582.

---

**Extended Data Fig. 1.**
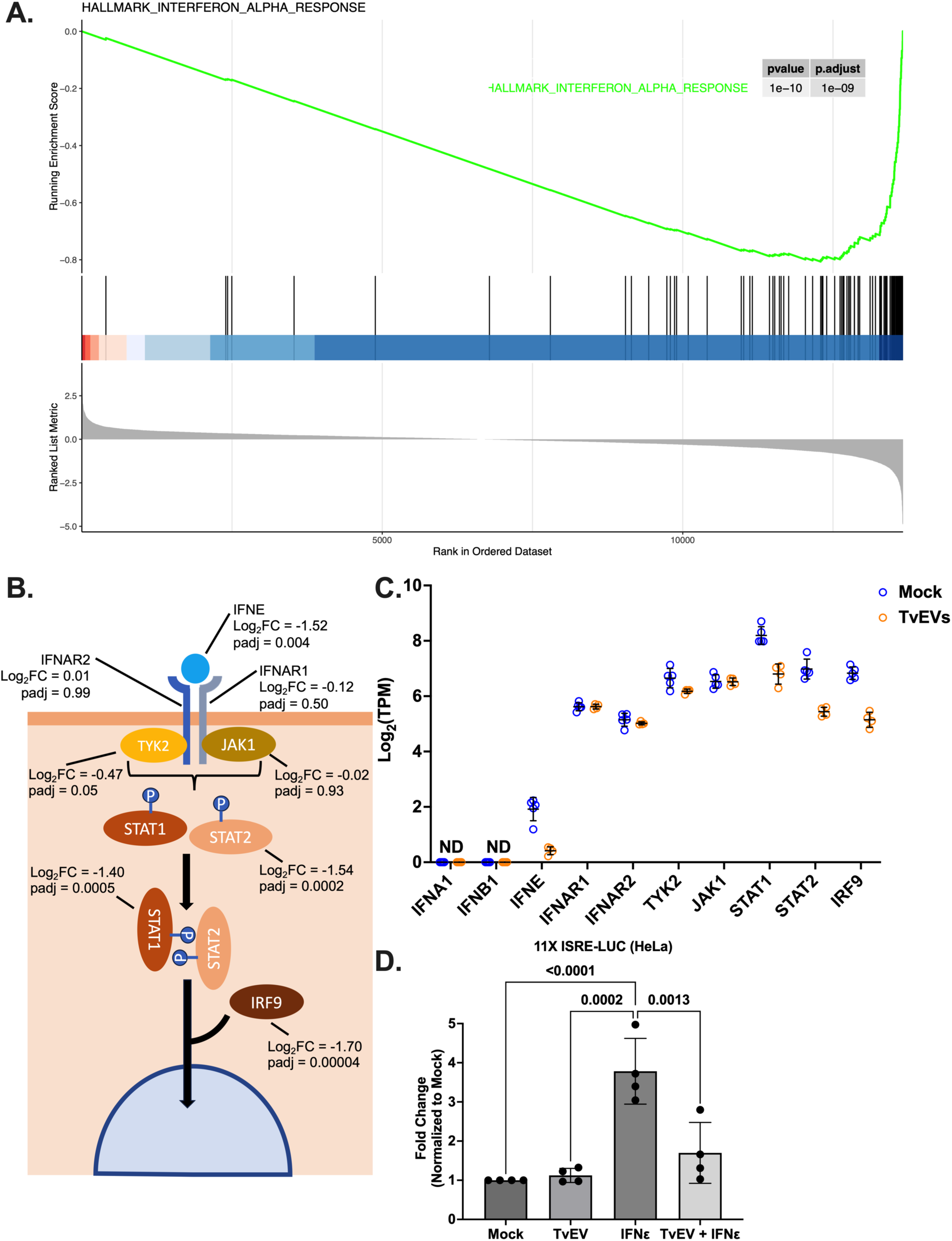
TvEVs downregulate the expression of multiple genes in the type I IFN pathway and suppress type I IFN responses. **a,** Gene set enrichment analysis (GSEA) showing Hallmark IFN alpha response of TvEV treated BPH-1 cells compared to mock treated BPH-1 cells. **b,** Simplified schematic of the type I IFN pathway: numbers represent p value of ≤ 0.05 and a log_2_ fold change for each indicated gene in the pathway from RNA-seq data in **Figure 2**. **c,** Transcript counts (represented in log_2_ transformation of Transcripts per million) for each individual gene in the type I IFN for Mock and TvEV treated BPH-1 cells from RNA-seq data in **Figure 2**. **d,** Luminescence is detected in HeLa cells expressing 11x ISRE Gaussia Luciferase reporter constructs treated with either TvEVs (10 µg/mL), IFNε (500 ng/mL), or TvEVs (10 µg/mL) + IFNε (500 ng/mL) for 6 hours. Luminescence is expressed as fold change ± SEM, compared to Mock. Bars, mean ± SD. N = 4 wells/experiment, 4 experiments total. Numbers above bars indicate p-values for one-way ANOVA, Dunnett’s multiple comparisons test.

**Extended Data Fig. 2.**
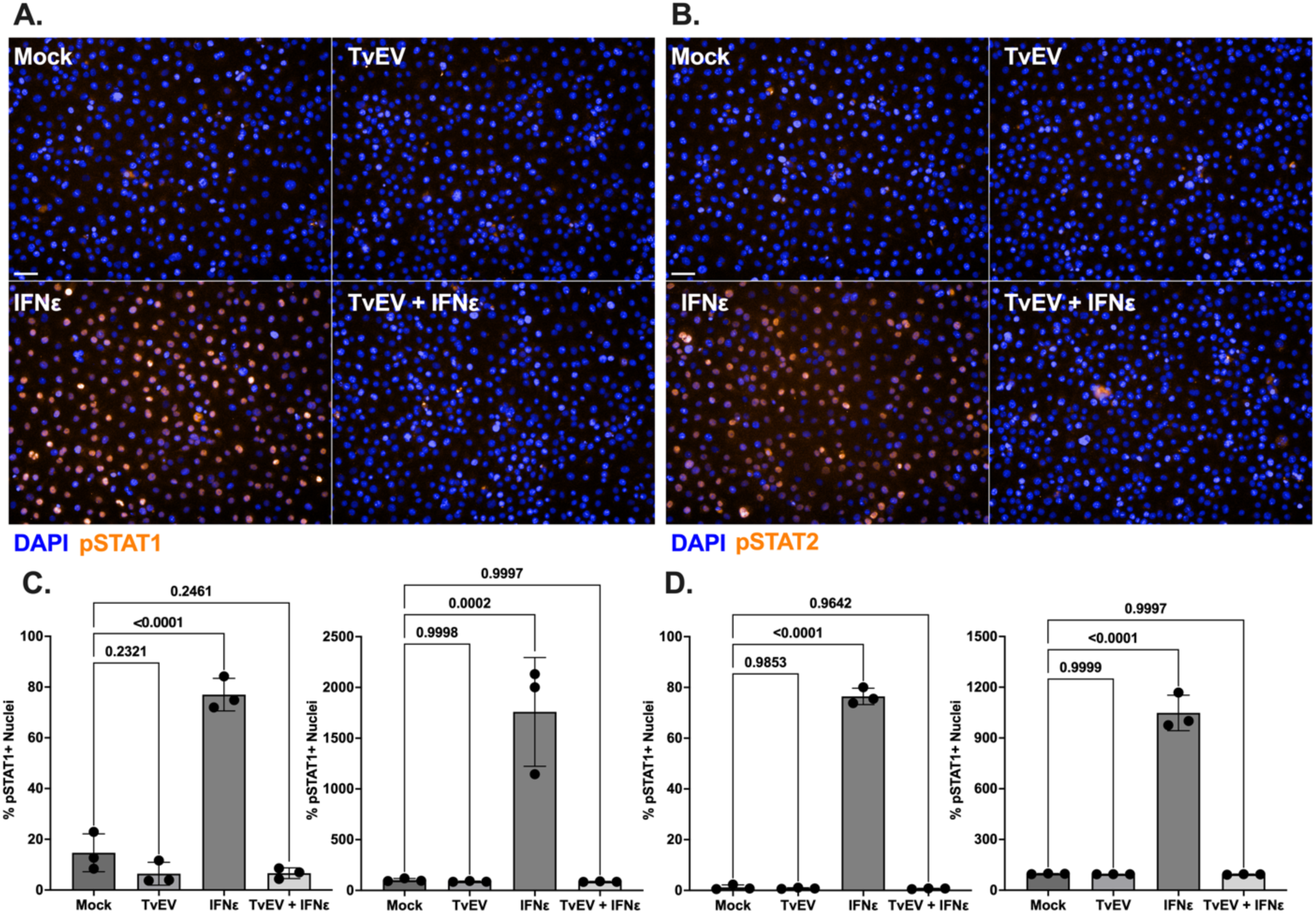
TvEVs block IFNε-mediated nuclear translocation of pSTAT1 and pSTAT2 in vaginal cells. **a and b,** pSTAT1/2 immunofluorescence in Ect1/E6E7 cells treated with either TvEVs (50 µg/mL), IFNε (500 ng/mL), or TvEVs (50 µg/mL) + IFNε (500 ng/mL). Depicted are DAPI staining Ect1/E6E7 and parasite nuclei (Blue) and pSTAT1/2 (Green). **c,** *Left,* quantification of percent of cells that at positive for nuclear pSTAT1. *Right,* Mean Fluorescent intensity (MFI) of nuclear pSTAT1. **d,** *Left*, quantification of percent of cells that at positive for nuclear pSTAT1. *Right*, Mean Fluorescent intensity (MFI) of nuclear pSTAT2. (**c** and **d**) Bars, mean ± SD. N = 3 wells/experiment, 9 fields of view/well, 3 experiments total. Numbers above bars indicate p-values for one-way ANOVA, Dunnett’s multiple comparisons test compared to vehicle control.

**Extended Data Fig. 3.**
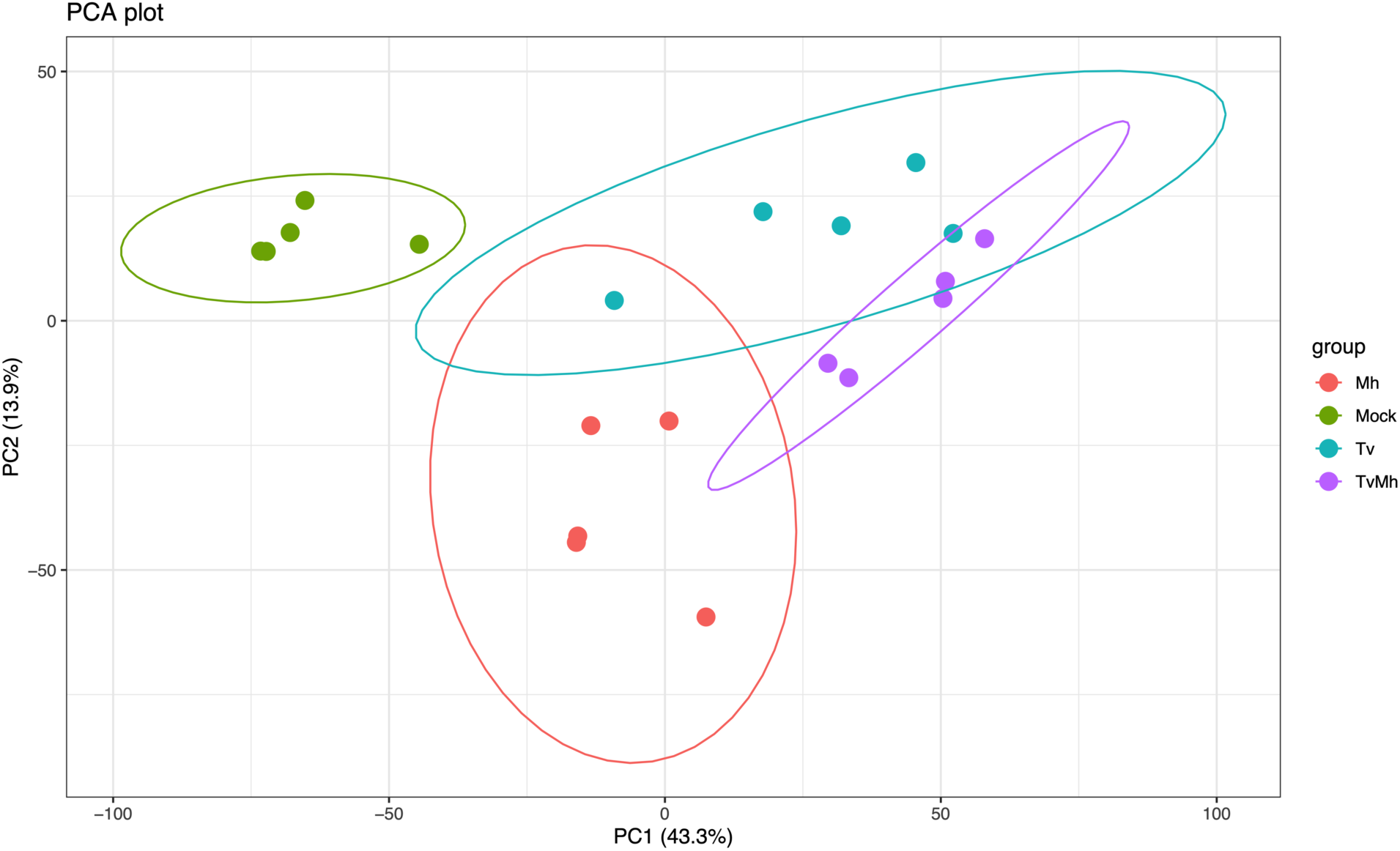
Principal component analysis of infected prostate cells. Principal components analysis showing distinct clustering of mock infected BPH-1 cells (Mock; *n* = 5), *T. vaginalis M. hominis* negative infected BPH-1 cells (Tv; *n* = 5), *M. hominis* infected BPH-1 cells (Mh; *n* = 5), and *T. vaginalis M. hominis* positive infected BPH-1 cells (TvMh; *n* = 5) from RNA-seq data generated in this study.

**Extended Data Fig. 4.**
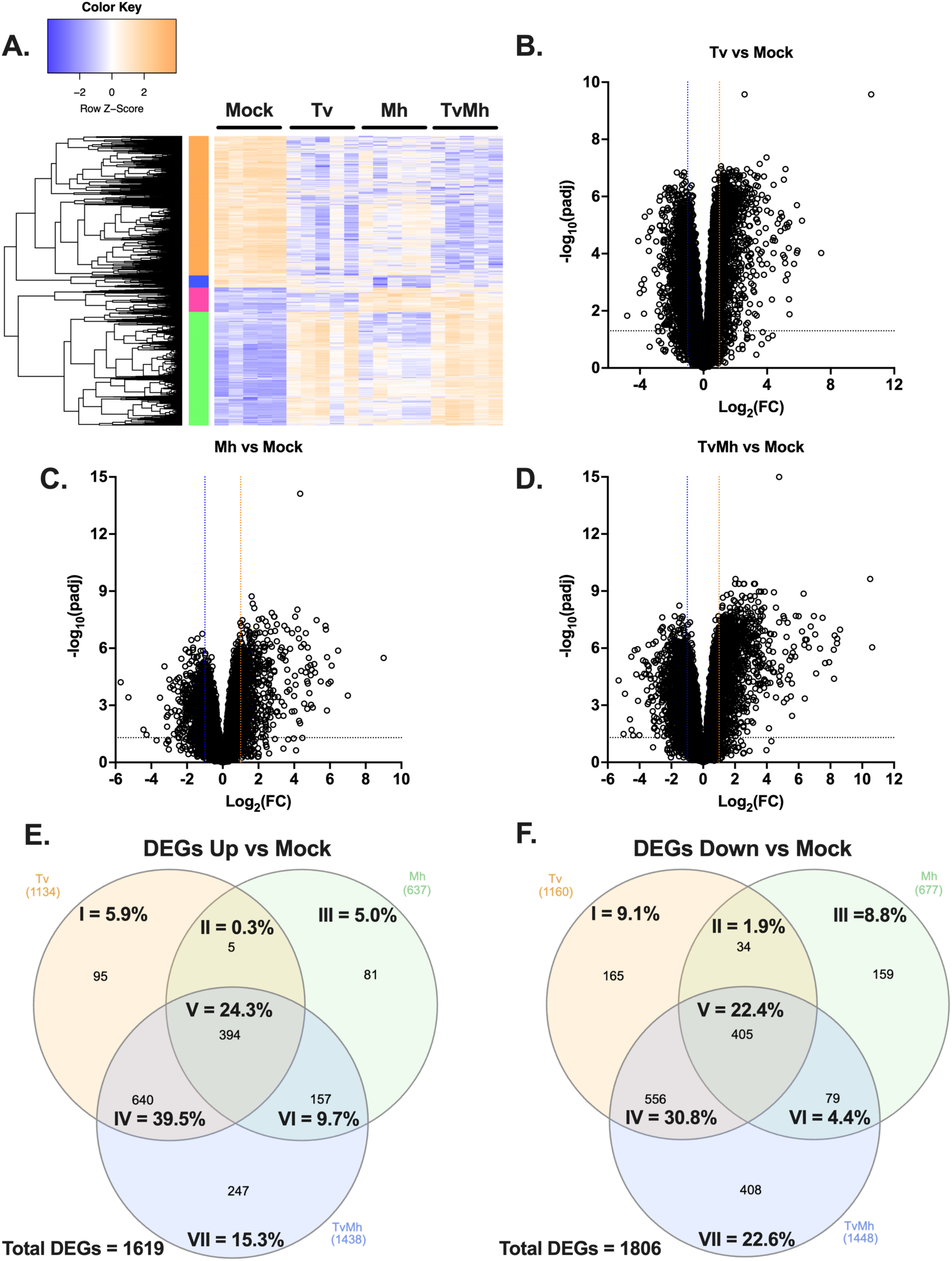
*T. vaginalis* and *M. hominis* elicit distinct and overlapping responses in host gene expression during infection. **a,** Heatmap of row z-score transformed genes identified as differentially expressed normalized to mock. **b-d,** Volcano plot depicting differences in gene expression between **b,** Tv infected cells compared to mock treated BPH-1 cells, **c,** Mh infected cells compared to mock treated BPH-1 cells, and **d,** TvMh infected cells compared to mock treated BPH-1 cells. The x axis corresponds to the log_2_ fold change in gene expression and the y axis indicates the adjusted p value. Genes with a −log_10_ p value of 1.3 or greater (p value of ≤ 0.05) and a log_2_ fold change −1 ≤ or ≥ 1 were deemed significantly differentially expressed. **e,** Venn diagram showing overlap in the number upregulated DEGs for each infection compared to mock control cells. **f,** Venn diagram showing overlap in of the number downregulated DEGs for each infection compared to mock control cells.

**Extended Data Fig. 5.**
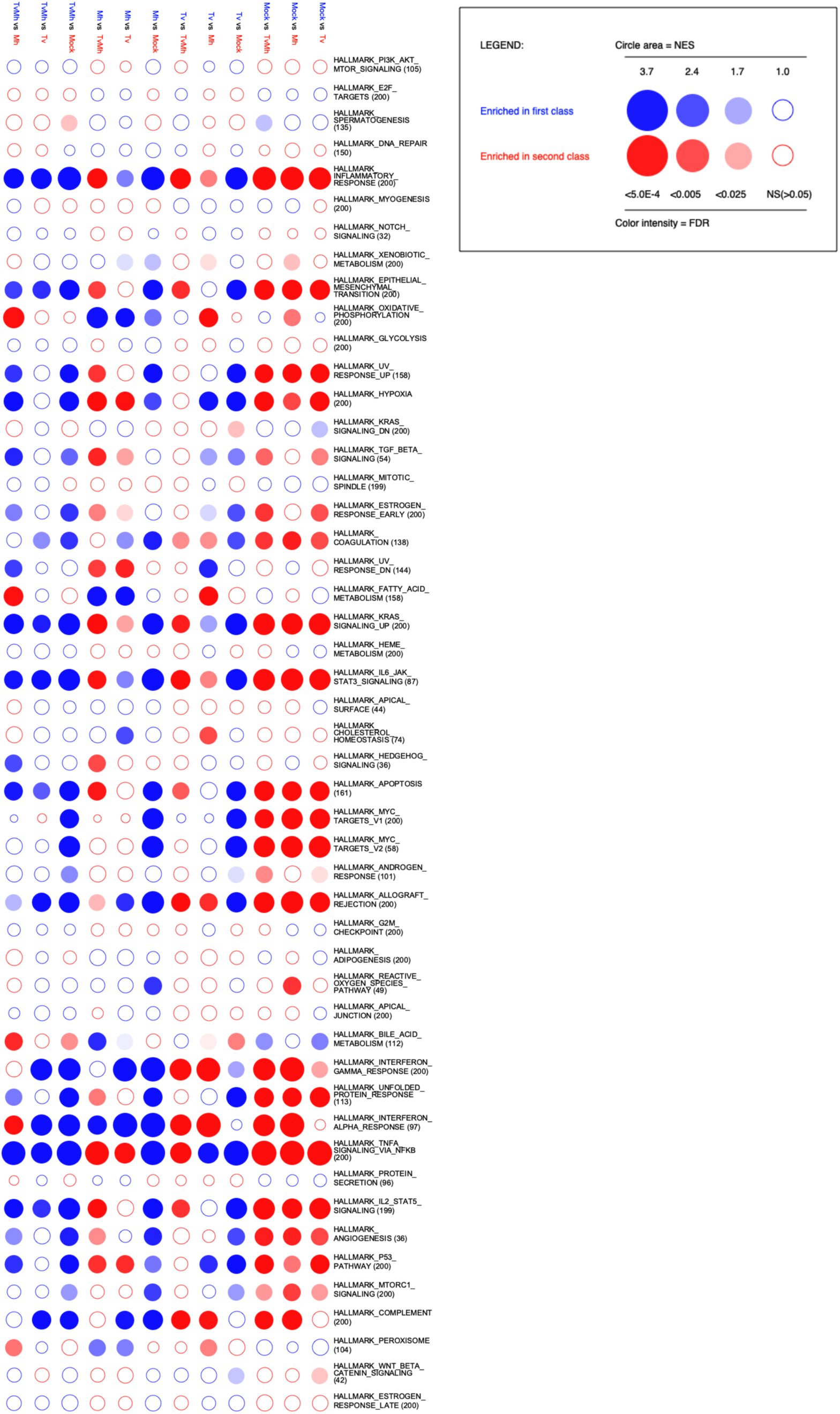
Extended GSEA Analysis of Tv, Mh, and TvMh infected prostate cells. Full BubbleMAP showing results of Gene Set Enrichment Analysis (GSEA) for the Hallmarks dataset for BPH-1 cells infected with Tv, Mh, or TvMh. Color and area of bubble represents normalized enrichment score (NES). Blue signifies pathway enriched in first class and Red signifies enrichment in second class. Intensity of color indicates false discovery rate (FDR).

**Extended Data Fig. 6.**
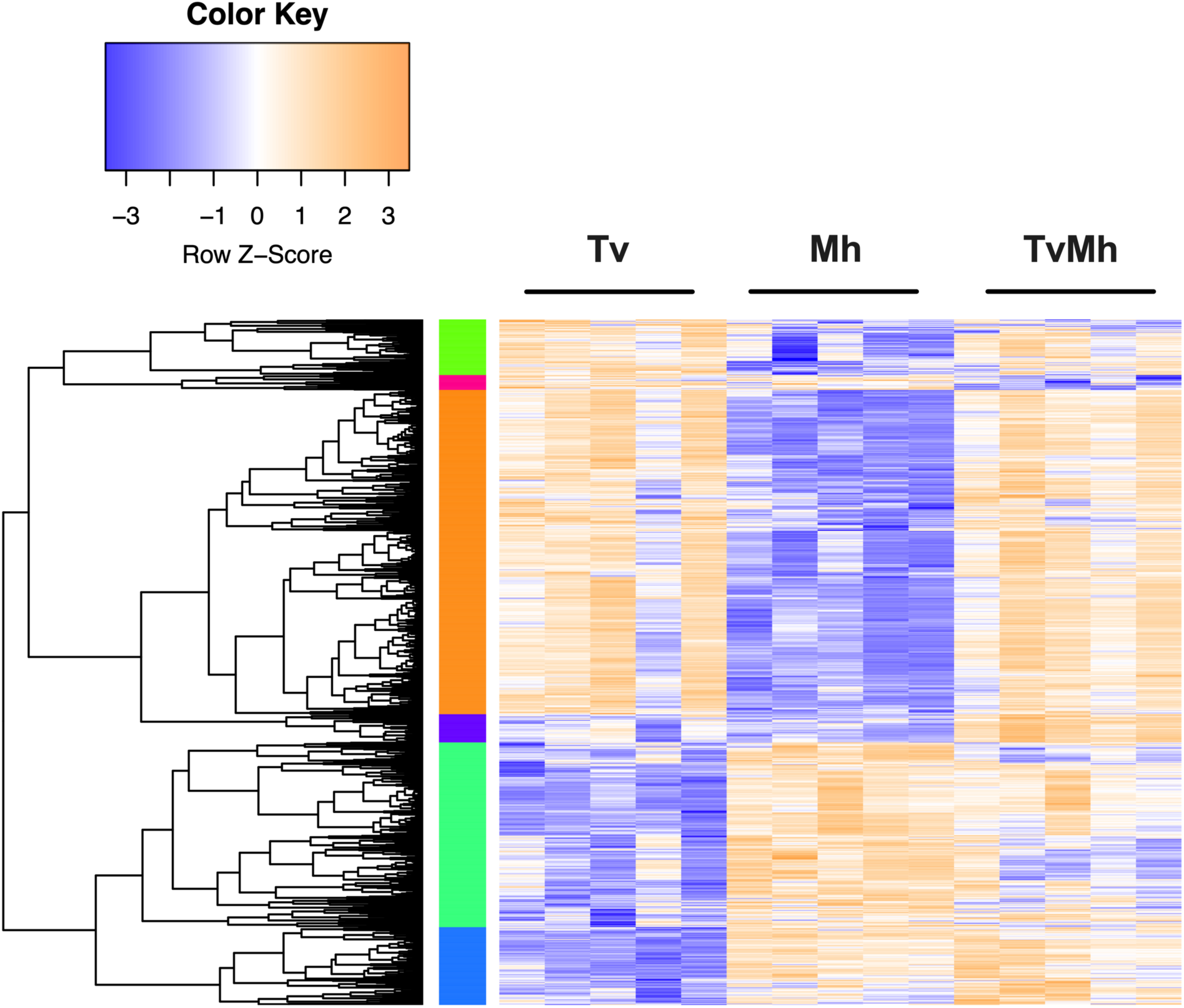
Heatmap of DEGs normalized to Tv infected BPH-1 cells. **b,** Heatmap of row z-score transformed 924 genes identified as differentially expressed (Log_2_(FC) ≥ 1 or Log_2_(FC) ≤ −1; padj ≤ 0.05) normalized to Tv infected BPH-1 cells.

**Extended Data Figure 7.**
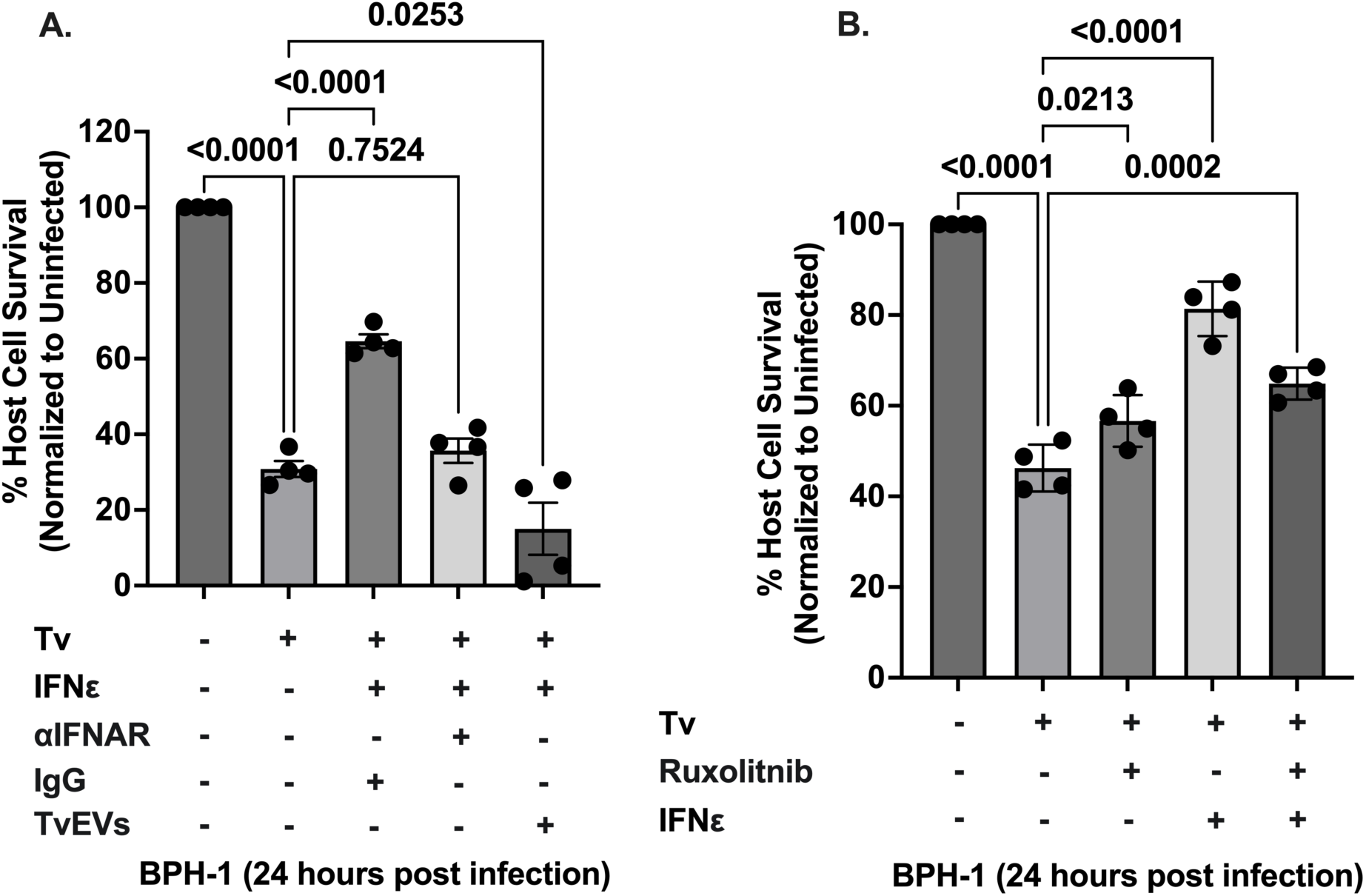
TvEVs and chemical disruption of the type I IFN pathway ablates IFNε-mediated protection against *T. vaginalis* cytolysis. **a** and **b,** Percent of host cell survival was quantified by enumerating the number of host cells left in the well after 24 hours compared to uninfected controls. Bars, mean ± SD. N = 3 wells/experiment, 51 fields of view/well, 4 experiments total. Numbers above bars indicate p-values for one-way ANOVA, Dunnett’s multiple comparisons test compared to uninfected control.

