## Supplemental_Information for "*Trichomonas vaginalis* extracellular vesicles suppress IFNε-mediated protection against host cell cytolysis"

### Figures and Figure Legends

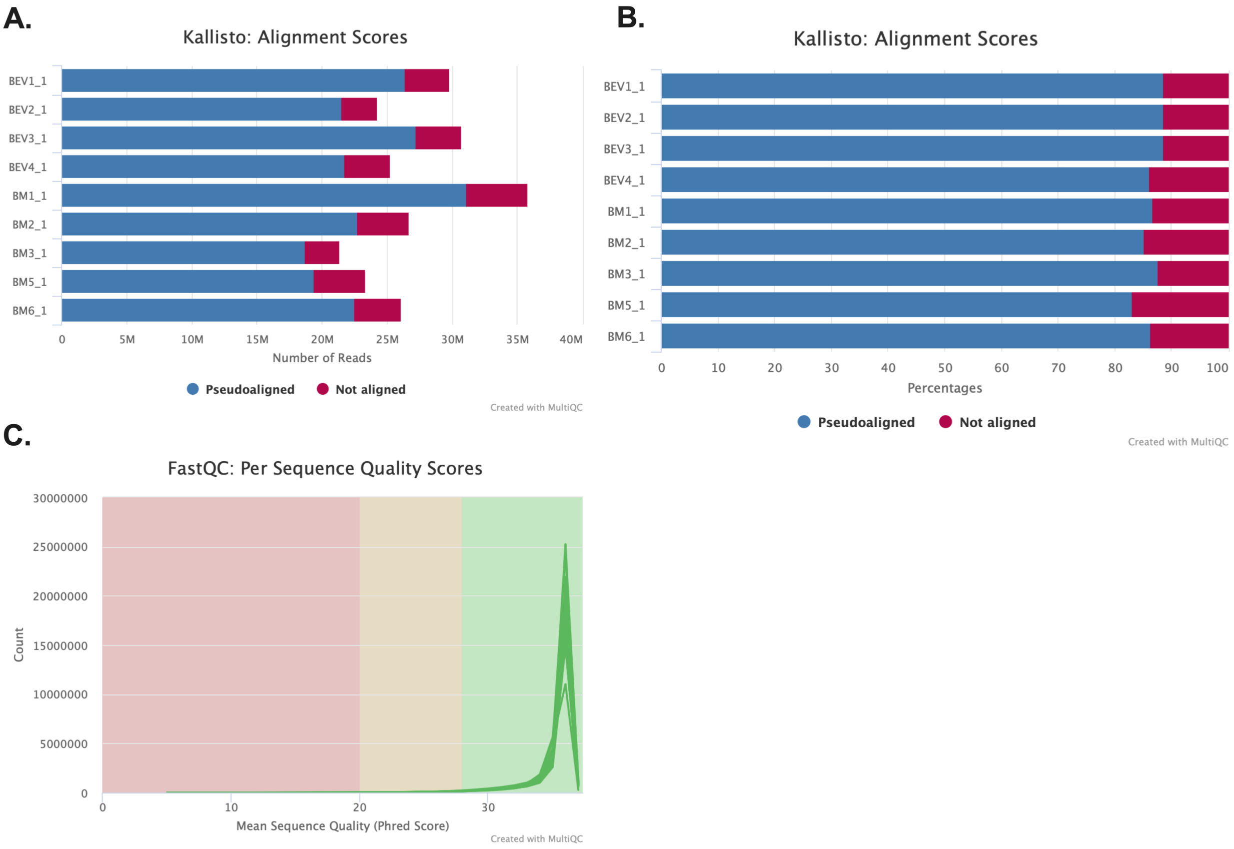


**Supplementary Fig. 1:** MultiQC summary of RNA-seq data in **Figure 2**. **a and b,** Summary of number of reads and percentage of reads for each sample that pseudoaligned to the human genome (EnsDb.Hsapiens.86). **c,** Per sequence Phred score for each sample summarized using MultiQC.

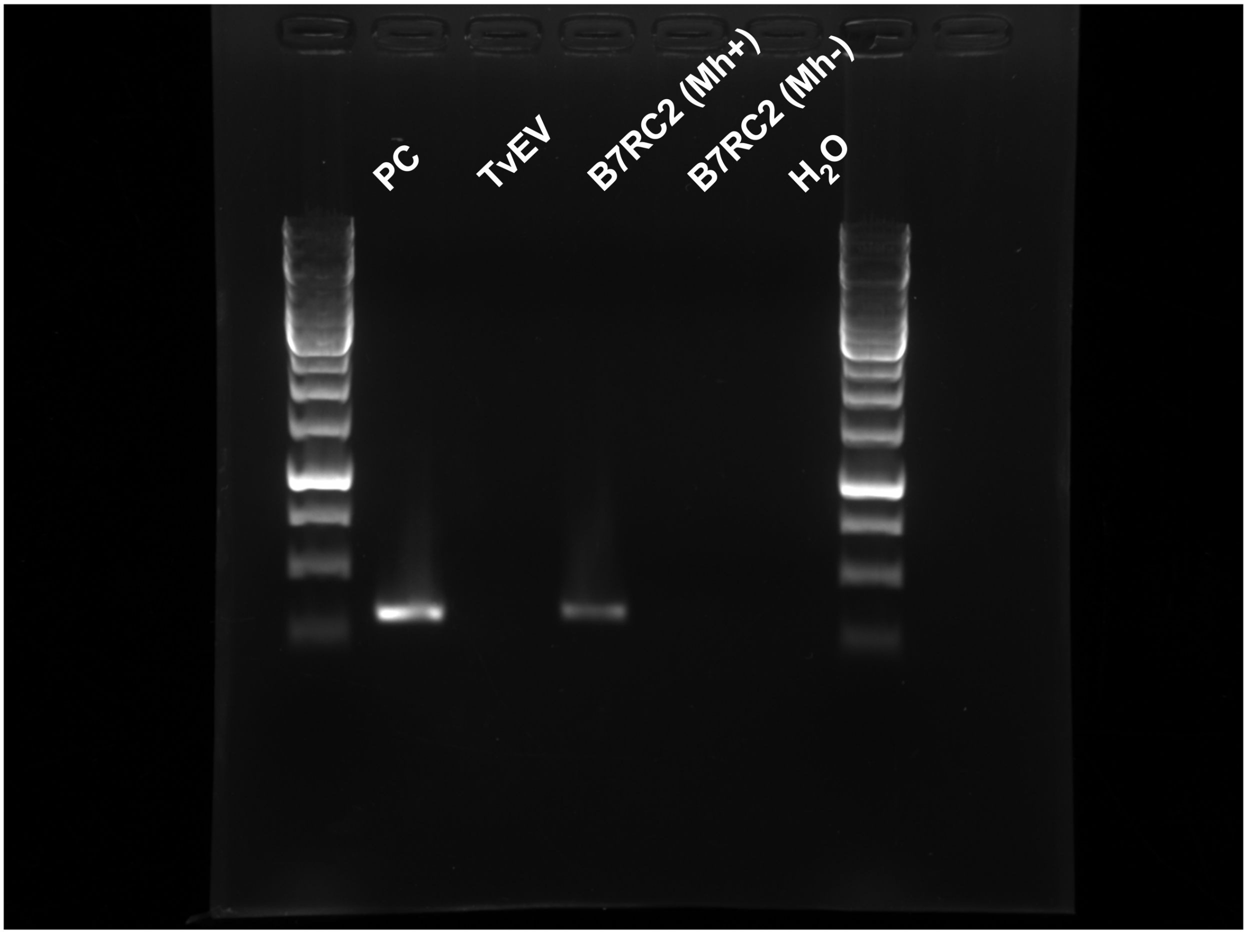


**Supplementary Fig. 2 PCR confirmation of *M. hominis* positivity:** PCR confirmation of *M. hominis* presence or absence using primers specific for *M. hominis* 16s RNA. Lane 1 DNA ladder, Lane 2 PC = positive control (*M. hominis* genomic DNA was used as a template), Lane 3 TvEV samples were confirmed clear of contaminating *M. hominis*, Lane 4 B7RC2 genomic DNA harboring *M. hominis* was used as a template, Lane 5 B7RC2 genomic DNA cleared of *M. hominis* was used as a template, Lane 6 $H_2O$ control, and Lane 7 DNA Ladder. Primers amplify expected band size of 334bp. Primers are 5'-CAATGGCTAATGCCGGATACGC-3' and 5'-GGTACCGTCAGTCTGCAAT-3'.

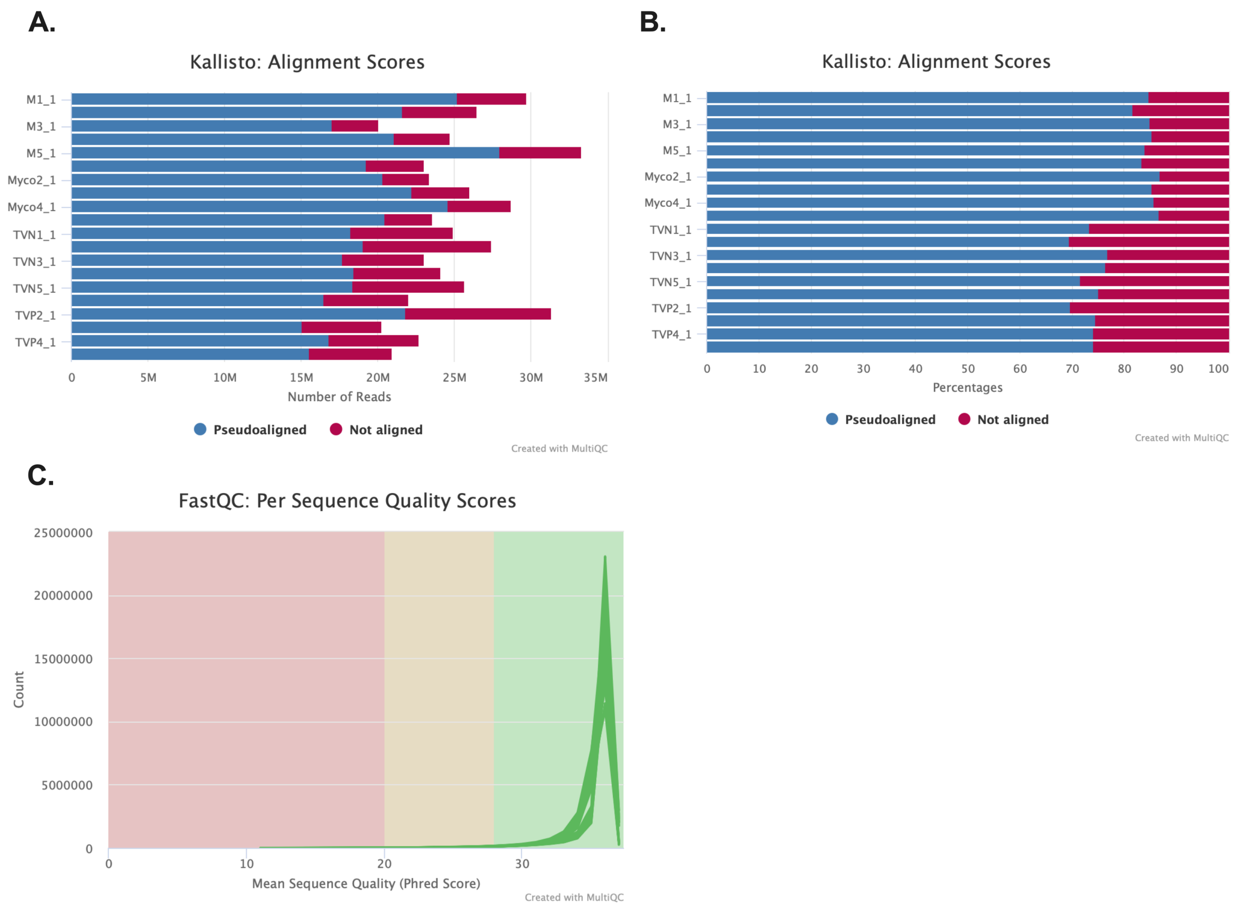


**Supplementary Fig. 3:** MultiQC summary of RNA-seq data in **Figure 4**. **a and b,** Summary of number of reads and percentage of reads for each sample that pseudoaligned to the human genome (EnsDb.Hsapiens.86). **c,** Per sequence Phred score for each sample summarized using MultiQC.

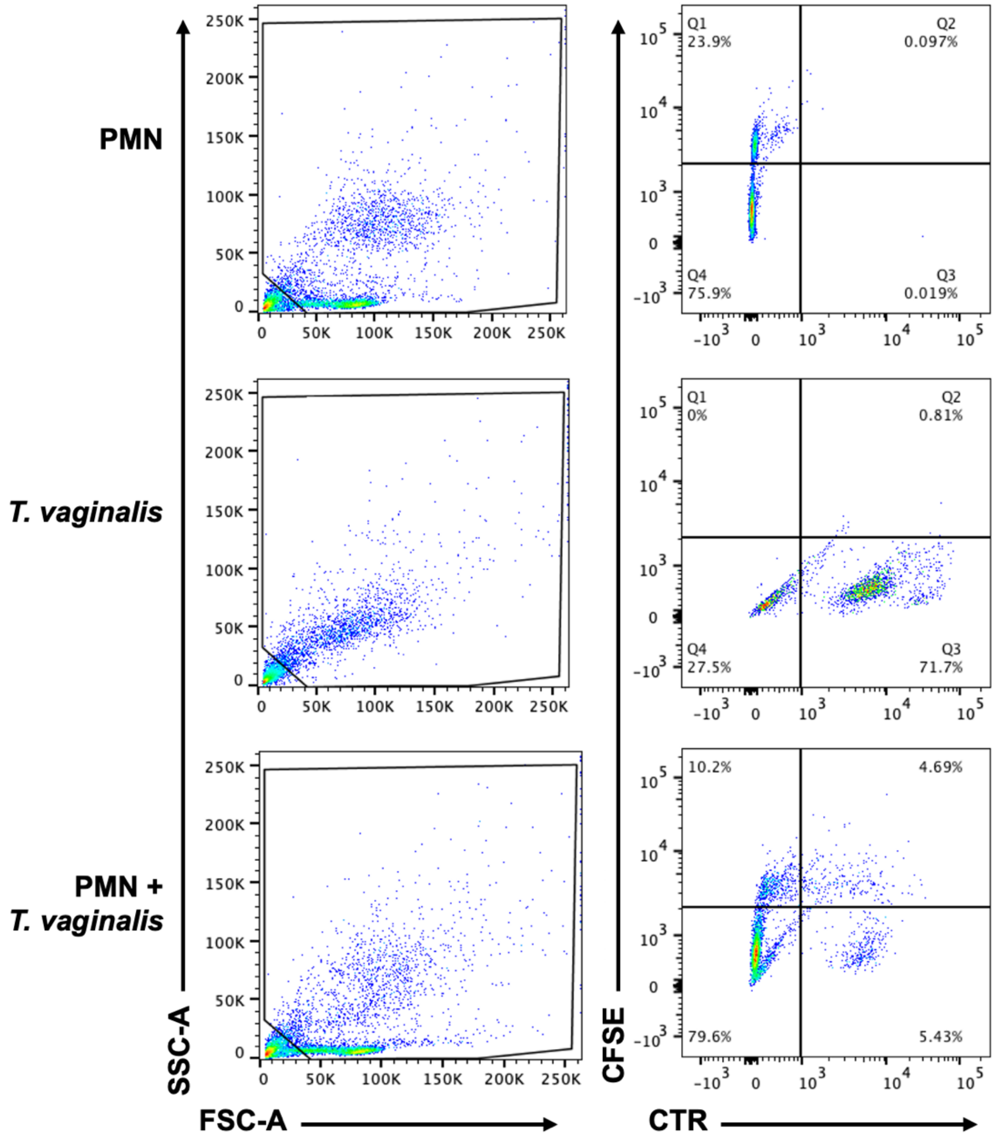


**Supplementary Fig. 4:** Gating strategy for flow cytometry analysis in **Fig. 6**. CFSE fluorescently stained polymorphonuclear cells (PMNs) were co-incubated with Cell Tracker Red (CTR) stained *T. vaginalis* at ratios of 16:1 and 8:1 and were left for 4 hours before being fixed with 4% paraformaldehyde. An initial forward and side scatter gating strategy excluded cell debris measuring below 50,000 and encompassed all the remaining PMN and *T. vaginalis*. Channels selecting CFSE and CTR were selected to quantify *T. vaginalis* compared to *T. vaginalis* alone samples. *Top row,* Gating strategy for PMN alone. *Middle row,* Gating strategy for *T. vaginalis*

alone. *Bottom row*, Gating strategy for PMN and *T. vaginalis* in co-culture. 10,000 total events were recorded for each condition.
